# Splenic granulopoiesis and S100A9 drive resistance to checkpoint inhibitors conferred by liver metastases

**DOI:** 10.1101/2025.09.30.679035

**Authors:** Rebecca J. Lee, Steven Hooper, Laura J. Pallett, Mariana Diniz, Tate McKinnon Snell, Zoe Ramsden, Nicolas Rabas, Stefan Boeing, Oliver Kennedy, Charlotte Buttercase, Gloryanne Aidoo-Micah, Gareth Price, Sarah C. Macfarlane, Sasha Bailey, Jessica Davies, Anandita Mathur, Ginevra Pistocchi, Alexandrine Carminati, Avinash Gupta, Patricio Serra, Brian Davidson, Joerg-Matthias Pollok, Venizelos Papayannopoulos, Ilaria Malanchi, Paul Lorigan, Mala K. Maini, Erik Sahai

## Abstract

Here, we investigate why liver metastases reduce the efficacy of immune checkpoint inhibitors (CPI). The poor prognosis of patients with liver metastases is associated with a systemic increase in neutrophils. Using experimental models, we confirm that mice with liver metastases respond poorly to CPI, have elevated neutrophils and suppress the response of subcutaneous lesions to CPI. We demonstrate that liver metastases, acting partly via IL-6, boost granulopoiesis in the spleen and promote the generation of immature S100A9^hi^ neutrophils that suppress T-cell proliferation. Human liver metastases exhibit a similar increase in S100A9^hi^ neutrophils. Neutrophil depletion attenuates the growth of liver metastases, but not subcutaneous metastases. Moreover, genetic deletion of S100A9 enables liver metastases to be effectively treated with CPI, and prevents liver metastases from suppressing the response in subcutaneous metastases. Thus, we document how liver metastases specifically change splenic granulopoiesis leading to changes in the microenvironment of non-hepatic lesions, and how targeting a key neutrophil protein restores the efficacy of CPI.

## Introduction

Immunotherapy has revolutionised the treatment of many cancers. Antibodies blocking immune checkpoints, specifically those targeting PD-1, PD-L1, CTLA4, or LAG3, are now used to treat an increasing number of cancer types as standard of care^1–4^. Dramatic improvements in survival have resulted, with cutaneous melanoma and microsatellite-instability–high colorectal cancer (CRC) being notable successes^2,4^. However, even in melanoma, 50-60% of patients do not get a lasting benefit from immune checkpoint inhibitors (CPI) targeting PD-1 alone or in combination with CTLA4 or LAG3 blockade^3,4^. Various features have been identified that distinguish patients who respond well to CPI compared to those who do not^5–7^. One of the key features associated with primary resistance to CPI is the presence of liver metastasis ^8,9^. This may reflect the tolerogenic function of the liver in certain physiological contexts; for example, to prevent excessive responses to factors derived from non-pathological gut microbiota or dietary antigens. The liver’s tolerogenic nature is exemplified by the fact that allogeneic liver grafts can be established and maintained without immunosuppression and that liver allo-transplantation can protect additional (non-liver) transplanted organs (from the same donor) from rejection^10,11^. A recent study showed that CD4^+^ T-regulatory cells (T_REG_) are activated in liver metastases, providing a potential explanation for the poor responses to CPI^12^. However, due to the complexity of this unique microenvironment, the drivers of CPI resistance in liver metastases are likely to be multi-faceted. A better understanding of the role of other suppressive cells within the tumour microenvironment, how they inter-relate and how liver metastases effect systemic immune response is required to overcome CPI resistance.

Neutrophils are an abundant myeloid cell type that can have both pro- and anti-tumorigenic properties^13,14^. Moreover, their heterogenous phenotype has been linked to differing responses to immunotherapy with Ly6E high, interferon-stimulated neutrophils associated with good outcomes, whereas FATP2 high, arginase producing, CCR5^+^ neutrophils are associated with resistance^15–18^. We have previously shown that immunosuppressive neutrophils can accumulate in the liver and inhibit antigen-specific T-cell responses in chronic hepatitis B infection^19^. These data suggest that neutrophils have a critical role in promoting immune tolerance in the liver microenvironment and could therefore be important in resistance to checkpoint inhibitors at this metastatic site.

In this study, we determine the mechanisms underlying the challenge of treating liver metastases with anti-PD-1 therapy, with a focus on melanoma. Through comparison of the same tumour cells growing within the liver parenchyma or at a subcutaneous location, both frequent sites of metastasis for cutaneous melanoma, we identify differences in neutrophil biology. Notably, liver metastases recruit pro-tumorigenic neutrophils, whereas subcutaneous lesions are associated with anti-tumorigenic neutrophils. We identify emergency granulopoiesis in the spleen as the source of immune suppressive neutrophils when liver metastases are present. Molecular profiling reveals these neutrophils to be an immature, ‘early neutrotime’ population of granulocytes with particularly high levels of S100A9. Moreover, genetic depletion of S100A9 reduces tumour growth and restores the sensitivity of liver metastases to anti-PD-1 therapy. In addition, through reducing systemic immune suppression, S100A9 depletion restores the responsiveness of subcutaneous metastases to CPI when liver metastases are present.

## Results

### Liver metastases are poorly responsive to immunotherapy with elevated neutrophils associated with inferior outcome

We first investigated the relationship between the presence or absence of liver metastases and outcomes in 1024 patients treated with immunotherapy. Consistent with previous reports, patients with advanced melanoma and non-small cell lung cancer (NSCLC) liver metastases had poorer overall survival (OS) compared to those without liver metastases following first line treatment with checkpoint inhibitors (melanoma n=641, NSCLC n=383, cohort characteristics Supp. Table 1). This difference was maintained even when adjusted for features associated with prognosis including age, sex, performance status, treatment type and for melanoma, BRAF status (Fig.1A median OS melanoma cohort 15.2 months liver metastases [95% confidence interval (CI) 11.6-24.5] vs. 44.9 months no liver metastases [95% CI 32.4-55.7] p<0.0001; and Supp.Fig.1A NSCLC cohort median OS 8.1 months liver metastases [95% CI 4.8-16.2] vs. 20.9 [95% CI 17.0-25.5] p=0.00022 Supp. Table2&3). Of note, disease progression in patients with melanoma liver metastases and extra-hepatic disease was not limited to the liver, but for the majority also occurred at other sites (n=62 Fig.1B), which supports pre-clinical data suggesting that liver metastases have a systemic impact^12,20^. Intrigued by previous data showing that liver metastases were associated with a reduced peripheral lymphocyte count and that high neutrophil to lymphocyte ratio (NLR) is significantly associated with poorer outcomes to CPI therapy, we examined these features in our cohort of patients with melanoma liver metastases^20–23^. We observed that in patients with melanoma liver metastases, a higher NLR was associated with poorer survival (median OS 12.7 months [95% CI 7.1-23.8] vs. 20.4 [95% CI 13.1-NA] p=0.001 Supp.Fig1B); however, there was no difference in survival when comparing high vs. low lymphocyte count (p=0.4, Supp.Fig1C). This indicated that changes in neutrophils drove the association between NLR and CPI outcomes. We therefore compared the levels of circulating neutrophils in a cohort of 108 patients with melanoma liver metastases who received either anti-PD-1 or combination anti-CTLA-4 plus anti-PD-1 therapy. Patients who progressed on immunotherapy had elevated levels of neutrophils compared to those who had a complete/partial response or stable disease for >6 months (Fig.1C). Moreover, Kaplan-Meier analysis indicated that neutrophil numbers equal or greater than the median were associated with shorter progression free (n=108, Fig.1D median 4.5 [95% CI 2.5-16.0] vs not reached [NR; CI 8.8-NR; p=0.0025) and overall survival (n=164, Fig.1E median 7.2 [95% CI 4.1-26.5] vs 21.9 [14.7-NR] months; p=0.007). These remained significant in Cox analysis of clinical prognostic factors in advanced melanoma, including sex, age, performance status, BRAF status, treatment type (combination/single agent) and baseline LDH (PFS HR 2.71 [95% CI 1.49-4.91], OS HR 2.56 [1.49-4.39]). Thus, we demonstrate that neutrophils are associated with reduced efficacy of CPI in human liver metastases.

**Figure 1.**
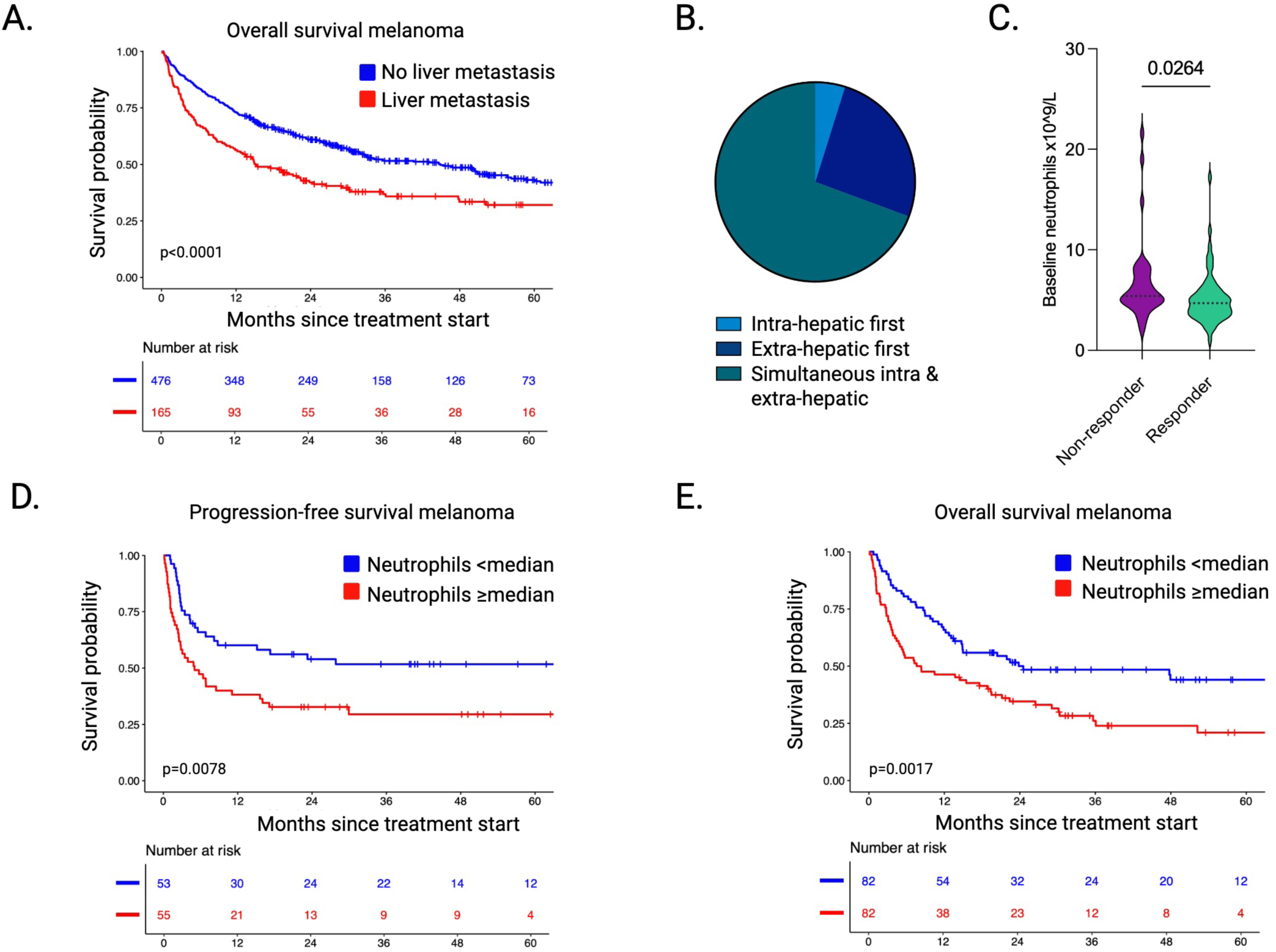
**Liver metastases are poorly responsive to immunotherapy with elevated systemic neutrophils associated with inferior outcome** **A.** Kaplan Meier survival estimate of patients with (n=165; red) or without (n=476; blue) melanoma liver metastasis treated with an anti-PD-1 containing regimen followed for a median of 48 months. Differences in survival determined by log-rank test. **B.** Distribution of different patterns of tumour progression in patients with intrahepatic and extrahepatic melanoma metastases present at baseline who progressed on checkpoint inhibitor therapy (CPI; n=62). **C.** Neutrophil count (x10^9^/L) prior to CPI in patients with melanoma liver metastasis who are non-responders vs. responders (defined as complete, partial and stable disease for >6 months) to anti-PD-1 containing CPI regimens. Differences between groups analysed using unpaired t-test. **D&E**. Kaplan Meier estimate of **D.** progression-free (n=108) and **E.** overall survival (n=164) in patients with melanoma liver metastases treated with an anti-PD-1 containing regimen stratified by systemic neutrophil count of the cohort ≥median (red) vs. <median (blue). Differences in survival determined by log-rank test.

### Liver metastases have a systemic impact on CPI efficacy

As melanoma frequently metastasises to the liver and to subcutaneous sites, we established an experimental model of liver metastases (LM) via ultrasound-guided orthotopic injection of syngeneic melanoma cells (4434 *BRAF* mutant cell line) and compared its behaviour to the same cells injected subcutaneously in the mouse flank (SC; Supp.Fig.2A). Mice with LM present treated with isotype control had significantly poorer survival compared to mice with SC only metastases (Fig.2A). The majority of mice with subcutaneous metastases had a complete response (CR) with anti-PD-1 treatment (17/19 mice; defined as no tumour measurable at experimental endpoint). In contrast, a minority of those with liver metastases also present had CR (7/35), with only a few days extension in survival attributable to CPI (Fig.2B, Supp.Fig.2B-H). The tumour growth rates of SC and liver lesions were similar, therefore the volume of disease could not account for the differential response (Supp. Fig.2I). We additionally observed that when 4434 melanoma cells were injected at both sites simultaneously, the presence of liver metastases prevented subcutaneous metastases from responding to anti-PD-1, resulting in decreased survival of the mice (Fig.2C, Supp.Fig.2D&G). Together, these data establish our preclinical model effectively recapitulates the resistance to CPI seen in patients with liver metastases. Furthermore, they indicate that liver metastases exert a systemic effect on lesions at an extrahepatic site.

**Figure 2.**
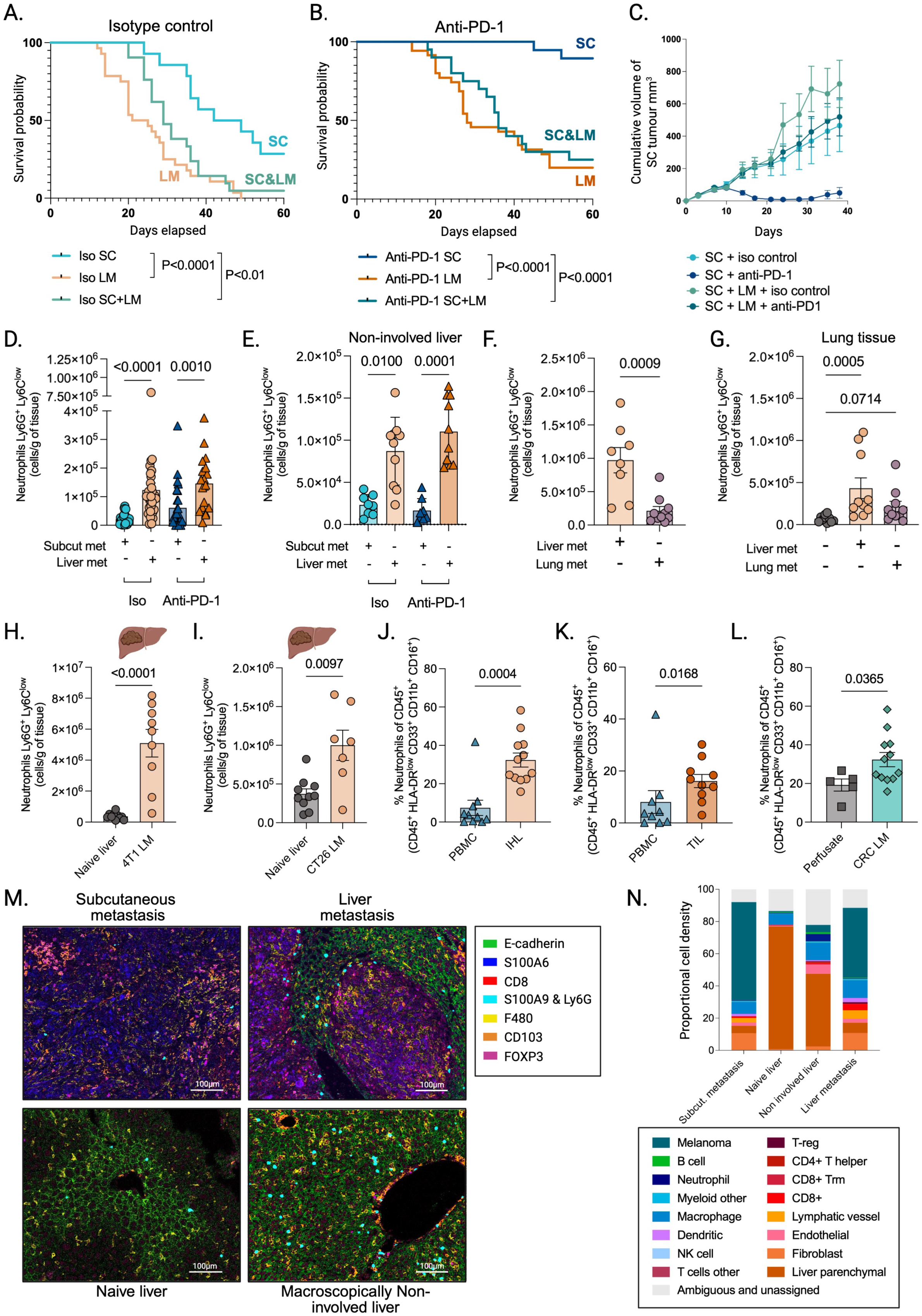
Liver metastases have a systemic impact on CPI efficacy and are associated with high frequencies of neutrophils both locally and systemically. **A&B.** Kaplan Meier survival estimate of mice (iso SC n=14, anti-PD-1 SC n=19, iso LM n=28, anti-PD-1 LM n=35, iso SC+LM n=21, anti-PD-1 SC+LM n=20) injected with 4434 melanoma cell line either subcutaneously (SC; blue) or intra-hepatically (LM; orange) or at both sites (SC+LM; green) and treated with **A** isotype control (iso; lighter shade) or **B** anti-PD1 (darker shades). Differences in survival determined by log-rank test. **C.** Cumulative growth (mm^3^) of subcutaneous metastases in animals with (green) or without (blue) liver metastases treated with isotype control or anti-PD-1 (n=7 biological replicates). **D&E.** Absolute count of Ly6G^+^Ly6C^low^ neutrophils present in **D.** metastasis and **E.** non-involved liver with subcutaneous (subcut; SC, blue) and liver metastases (LM, orange) present treated with an isotype control or anti-PD-1 (n=6 experiments; n=23 iso SC, n=31 iso LM, n=28 anti-PD-1 SC, n=18 anti-PD-1 LM) SC. **F.** Absolute count of Ly6G^+^Ly6C^low^ neutrophils in lung (generated by tail vein injection) and liver metastases (n=2 experiments). **G.** Absolute count of Ly6G^+^Ly6C^low^ neutrophils present in lung tissue in the naïve lung or when lung or liver metastases present (n=2 experiments). **H&I.** Absolute count of Ly6G^+^Ly6C^low^ neutrophils present in liver metastases compared to naïve liver (n=2 experiments) 14 days after injection of **H.** 4T1 breast cancer cells or **I.** CT26 colorectal cancer. **J&K.** Frequency of human HLA-DR^low^CD33^+^CD11b^+^CD16^+^ low density neutrophils (as a proportion of live CD45^+^ leukocytes) in **J.** intrahepatic leukocytes (IHL) and peripheral blood (PBMC) or **K.** tumour infiltrating lymphocytes (TIL) and PBMC of patients undergoing surgical resection for colorectal cancer (CRC) liver metastases. **L.** Frequency of HLA-DR^low^CD33^+^CD11b^+^CD16^+^ neutrophils (as a proportion of live CD45^+^ leukocytes) in IHL of patients with CRC liver metastases compared to IHL obtained by perfusion of healthy donor livers prior to solid organ transplantation. **M.** Representative image obtained by imaging mass cytometry of a subcutaneous metastasis, liver metastasis, naïve liver and non-involved liver of animals with LM. Cell types: Liver stroma (Ecadherin^+^, green), melanoma (S100A6^+^, dark blue), CD8^+^ T-cells (CD8^+^, red) neutrophils (S100A9^+^ or Ly6G^+^, light blue) macrophages (F480^+^, yellow) dendritic cells (CD103^+^, orange) T-regulatory cells (FOXP3^+^, pink). Scale bar 100 µm. **W.** Stacked bar chart showing the proportions of different cell types within different tissues identified using imaging mass cytometry. Deep learning-based cell segmentation (deep-imcyto) and automated cell-type annotation (TYPEx) using the TRACERx-PHLEX pipeline were used to identify the different cell types. Subcutaneous metastasis (n=24 tissue micro-arrays [TMA]), liver metastasis (average of n=13 TMA), naïve liver (n=4 TMA) and non-involved liver (LM present n=8 TMA). Proportional cell density = raw count/total raw count of sample. All error bars represent mean ±S.E.M. Mann-Whitney test used for comparison of 2 groups (F, H-L). Kruskal-Wallis test used for comparison of >2 groups (D, E, G).

### The immune environment of liver metastases

Motivated by our analysis of patients, we investigated the immune microenvironment of murine liver metastases compared to macroscopically non-involved liver (termed “non-involved”), naïve liver (tumour-free mice), and subcutaneous lesions. There was a clear increase in the number of Ly6G^+^ Ly6C^low^ neutrophils in the liver compared to subcutaneous metastases with and without anti-PD-1 treatment (Fig.2D, gating strategy Supp.Fig.2J). Of note, the increase in neutrophils was observed throughout the non-involved liver, suggesting a global change in the liver microenvironment, and possibly systemically, when a liver metastasis is present (Fig.2E). Natural killer (NK) cell numbers were not different between sites (Supp.Fig.2K). The number of CD8^+^ T cells was marginally lower in the liver, although this difference was not significant in anti-PD-1 treated animals (Supp.Fig.2L). The proportion of effector memory CD44^+^ CD62L^-^ CD8^+^ T cells was lower in liver metastases following anti-PD-1 treatment compared to subcutaneous disease (Supp.Fig.2M). T_REG_ numbers were similar between liver and subcutaneous disease, with no consistent phenotypic changes such as CD69, ICOS or CTLA4 expression that would indicate an increase in their immune suppressive capacity (Supp.Fig.2N-Q).

Next, we assessed whether neutrophil numbers were consistently higher in the liver compared to other metastatic sites. This revealed that there were increased neutrophils in liver metastases compared to lung metastases (Fig.2F). Further, lung metastases did not have significantly elevated neutrophil numbers compared to the lungs of tumour-free (naïve) mice, whereas an increase was observed in the lungs of mice with liver metastases present (Fig.2G). We further validated the high numbers of neutrophils present in liver metastases and in the non-involved liver in models of breast cancer (4T1) and CRC (CT26, Fig.2H&I and Supp.Fig.2R&S).

Consistent with these data, the levels of immature, low density neutrophils (CD45^+^, HLA-DR^lo^, CD33^+^, CD11b^+^, CD16^+^; gating strategy Supp.Fig.2T) in the liver and liver metastases from patients with CRC were higher in both the intrahepatic lymphocyte compartment (IHL) and in tumour-infiltrating lymphocytes (TIL) compared to the periphery (Fig.2J&K – we analysed CRC metastases as these are more commonly resected than melanoma liver metastases). Critically, we observed higher numbers of neutrophils in livers with CRC liver metastases compared to perfusates taken from healthy donor transplant livers, suggesting the presence of liver metastasis is associated with hepatic neutrophil accumulation (Fig.2L). Together, these data indicate that liver metastases lead to increases in neutrophil numbers and that these increases are not restricted to the liver lesion, but are observed in non-involved liver and lung tissue.

The functionality of immune responses depends on the location and proximity of different cell types; therefore, we performed highly multiplexed immunostaining of naïve liver, macroscopically non-involved liver (with a liver metastasis present), liver metastases, and subcutaneous melanoma metastases. This confirmed the increase in neutrophils when liver metastases were present compared to naive liver, with the majority of neutrophils at the tumour edge and in the liver paraenchyma (Fig.2M&N). Neighbourhood analysis identified four predominant types of cellular landscape, with cluster 1 predominantly melanoma, cluster 2 dominated by leukocytes but also containing melanoma cells, cluster 3 containing fibroblasts and cluster 4 mainly liver parenchyma (Supp.Fig.2U). Neutrophils were predominantly in cluster 2 & cluster 4 neighbourhoods located in tumours and the liver parenchyma, respectively (Supp.Fig.2U&V). Of note, neutrophils were co-located with CD8^+^ T cells and T_REG_ around the margins of liver metastases (Fig.2M, Supp.Fig.2V-X). The presence of a liver metastasis also increased cluster 2 neighbourhoods in the non-involved liver, with considerably more diverse immune infiltrates apparent compared to naïve liver (Supp.Fig.2V-X). CD8^+^ T cells were able to effectively infiltrate liver metastases, indicating that liver lesions are not ‘immune cold’ or ‘excluded’ (representative images Fig.2M with quantification in Fig.2N). In subcutaneous metastases, neutrophil cell neighbourhoods contained more melanoma cells than in liver metastases, whereas there were more T_REG_ present in neutrophil neighbourhoods in liver metastases (Supp.Fig.2X). Overall the cellular composition of neighbourhoods was largely similar between liver and subcutaneous metastases (Supp.Fig.2W), suggesting that qualitative differences in cell states and cell interactions, rather than fundamental spatial differences, are more likely to underpin differential responses to checkpoint therapy.

### Neutrophils have divergent functions in liver and subcutaneous lesions

Previous studies have shown that neutrophils can promote tumour growth and metastasis^24,25^. The data described above raise the possibility that neutrophils may be responsible for the divergent response of liver and subcutaneous metastases to CPI. To test this, we depleted neutrophils using a Gr-1 antibody, with monocytic cell numbers unaffected (Supp.Fig.3A&B). Intriguingly, this had differing effects on liver and subcutaneous melanoma metastases. In the context of liver metastases, targeting Gr-1 led to an extended survival, while the survival of mice with subcutaneous metastases was reduced by anti-Gr-1 antibodies (Fig.3A&B). We confirmed the improved survival of mice with liver metastases upon neutrophil depletion using a more specific anti-Ly6G antibody (clone 1A8, Supp.Fig.3C). We hypothesised that the pro-tumorigenic function of neutrophils in liver metastases may relate to their ability to suppress T cells. In support of this, neutrophils purified from liver metastases or from non-involved liver tissue of a mouse with a liver metastasis suppressed the *in vitro* proliferation of their CD3/CD28-stimulated T cells much more effectively than neutrophils from the livers of tumour-free (naïve) mice (Fig.3C). Together, these data demonstrate that immuno-suppressive neutrophils promote the growth of liver metastases but can act to restrain subcutaneous lesions if there is no liver metastasis present. Combined treatment with anti-Gr-1 and anti-PD-1 led to an additive increase in survival in mice with liver metastases compared to untreated mice than either treatment given alone. Intriguingly, the negative effect of neutrophil depletion was no longer observed in the subcutaneous metastasis when treated with anti-Gr-1 plus anti-PD-1, suggesting a dominant effect of anti-PD-1 in this context (Fig.3A&B). Thus, neutrophils are associated with different effects on tumour growth and mouse survival, depending on site of metastasis and treatment.

**Figure 3.**
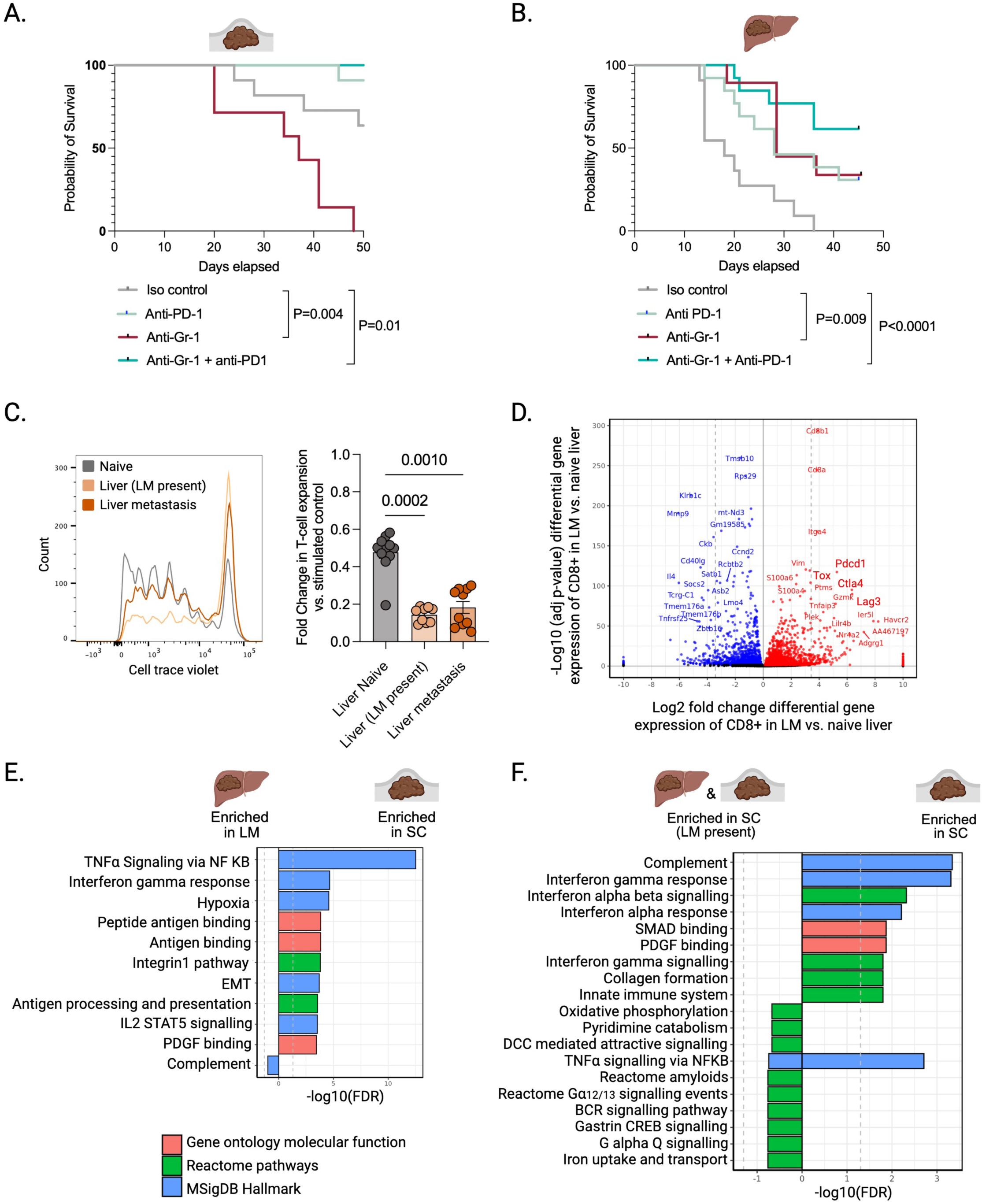
Depletion of neutrophils has a differential effect on tumour growth in the liver compared to subcutaneous site. **A&B.** Kaplan Meier survival estimate of mice injected **A.** subcutaneously or **B.** intrahepatically with 4434 melanoma cell line and treated with isotype control (grey), anti-PD-1 (light green), anti-Gr-1 (red) or anti-PD-1 plus anti-Gr-1 (teal). Differences in survival determined by log-rank test. **C.** Fold change in T cell expansion (from naïve mice) as measured by cell trace violet dilution after CD3/CD28 bead stimulation following 5-7 days of co-culture 1:1 with neutrophils isolated from either naïve livers, LM or non-involved liver with a LM present compared to culture alone. Summary data of n=3 experiments shown with a representative cell-trace violet plot. Kruskal-Wallis test performed to compare groups. **D.** Volcano plot depicting differential gene expression analysis of single cells in the CD8^+^ cluster infiltrating liver metastasis compared to naïve liver (red=increased, blue=decreased, black=unchanged expression, n=2 experiments). **E&F.** Hallmark pathway analysis examining differentially expressed genes enriched in single cell RNA sequencing analysis of facs-sorted neutrophils infiltrating **E.** subcutaneous vs. liver metastases **F.** subcutaneous metastases with vs. without liver metastases present (n=2 experiments, 1 biological replicate per experiment.

To obtain a deeper understanding of differences in immune cell state between liver and subcutaneous lesions, we performed single-cell RNA sequencing (scRNA-seq) on tumours two weeks after the injection of tumour cells. This revealed that CD8^+^ cells in liver metastases had a more exhausted phenotype compared to CD8^+^ cells in naïve liver, as evidenced by high Lag3, Pdcd1 (PD-1), Ctla4 and Tox gene expression (Fig.3D). Critically, neutrophils had a heterogenous phenotype depending on the infiltrating site, with IFNγ response and TNFα signalling pathways more highly expressed by neutrophils in the subcutaneous metastasis compared to the liver metastasis (Fig.3E). Furthermore, the presence of liver metastasis suppressed interferon gamma signalling in neutrophils infiltrating subcutaneous lesions (Fig.3F). Together, these data indicate that liver metastases have a systemic impact on both neutrophil state and function.

### Liver metastases drive emergency granulopoiesis in the spleen

Having established the importance of neutrophils for tumour promotion in the context of liver metastases, we sought to determine their origin. To this end, we pulsed either tumour-free (naïve) mice or mice harbouring subcutaneous or liver melanoma metastases with 5-ethynyl-2’-deoxyuridine (EdU) for one hour before analysing neutrophils in the non-involved liver, at the metastatic sites, blood, bone marrow and spleen (Fig4.A-E, gating Supp.Fig.4A). There was little neutrophil progenitor proliferation in the liver tissue itself or in the blood in any condition analysed. Although, the bone marrow was the major site of neutrophil production in mice without tumours, neither the presence of liver nor subcutaneous metastases had any impact on neutrophil progenitor proliferation at that site. In contrast, the presence of liver, but not subcutaneous, metastases boosted the neutrophil progenitor proliferation in the spleen. This increase in proliferating splenic neutrophils was independent of the size of liver metastasis (Supp.Fig.4B). We also assessed whether this was a response to the liver wounding caused by the injection, but did not observe any increase of neutrophils in the spleen of mice injected with Matrigel alone compared to mice with no injection (Supp.Fig.4C). We further validated these findings in the 4T1 and CT26 models which also showed a significant increase in number of neutrophils in the spleens of mice with liver metastases present compared to those without (Supp.Fig.4D&E). To assess whether there could be specific liver-spleen mediators of communication, we assess cytokine/chemokine lysates of liver metastases compared to subcutaneous metastases and observed higher levels of IL-6 in the liver metastases (Supp.Fig.4F), with scRNA seq data indicating Kupffer cells as a source (Supp.Fig.4G). Treatment with anti-IL-6R reduced the numbers of neutrophils in the spleens of mice bearing liver metastasis (Supp.Fig.4H). Thus, the presence of metastases in the liver specifically drives splenic production of neutrophils and has a more limited effect on granulopoiesis in the bone marrow.

**Figure 4.**
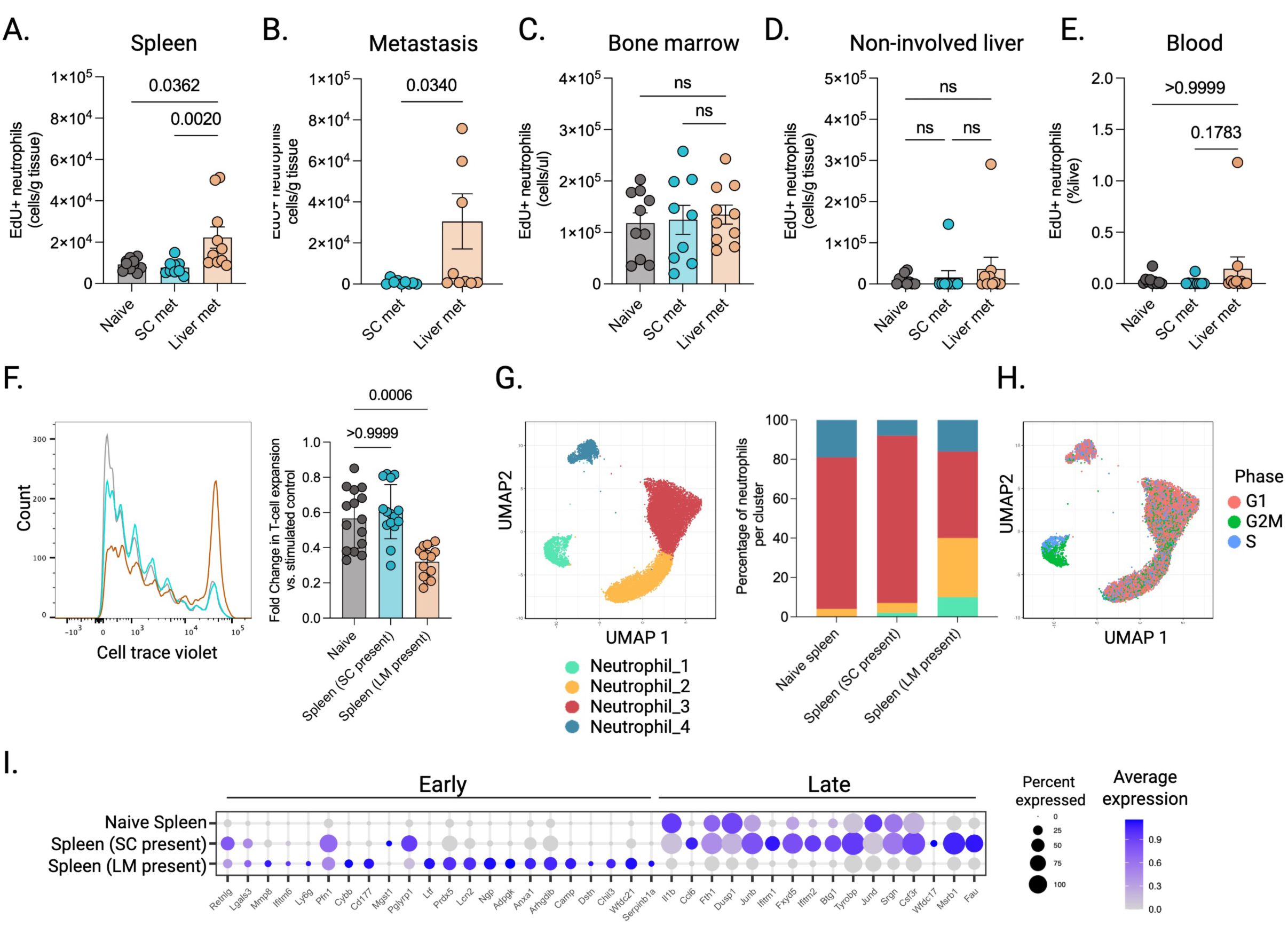
Neutrophils proliferate in the spleen and have a more immunosuppressive phenotype when a liver metastasis is present. **A-E.** Absolute count of 5-ethynyl 2’-deoxyuridine (EdU) positive Ly6G^+^Ly6C^low^ neutrophils in the **A**. spleen **B**. metastasis **C**. bone marrow **D**. non-involved liver and **E.** blood in naïve mice (naïve; grey) vs. mice with a subcutaneous (SC met; blue) or liver metastasis present (LM; orange) n=2 experiments. Mann-Whitney test used for comparison of 2 groups. Kruskal-Wallis test used for comparison of 3 groups. **F.** Fold change in T cell expansion (from naïve mice) as measured by cell trace violet dilution after CD3/CD28 bead stimulation following 5-7 days of co-culture 1:1 with neutrophils isolated from either naïve spleens or spleens with a SC or LM present compared to culture alone. Summary data of n=3 experiments shown with a representative cell-trace violet plot. **G.** Single cell RNA sequencing UMAP plot of splenic neutrophils divided into 4 clusters (1-4) with summary bar plot showing proportion of neutrophils in each cluster within naïve spleens or spleens of mice bearing subcutaneous or liver metastases (n=5 biological replicates per group). **H.** UMAP visualisation showing single neutrophils which are in G2M or S phase depending on neutrophil cluster (n=5 biological replicates per group). **I.** Single cell analysis comparing gene expression of early and late neutrotime signatures of neutrophils in naive spleens or spleens of mice with a subcutaneous or liver metastasis present (n=5 biological replicates per group). Dot size represents percentage of cells expressing and colour represents average gene expression. All error bars represent mean ±S.E.M.

Next, we sought to understand if the neutrophils produced in the spleen in response to liver metastases were functionally distinct and what might be mediating changes in splenic granulopoiesis. We observed that the presence of liver, but not subcutaneous, metastases increased the ability of splenic neutrophils to suppress T-cell proliferation in response to anti-CD3/CD28 engagement (Fig.4F). In addition, following TCR engagement, CD8^+^ T cells produced less TNFα and exhibited a reduced capacity to mobilise cytotoxic granules to their surface after co-culture with splenic neutrophils extracted from mice with liver metastases compared to those without (Supp.Fig.4I&J). Interestingly, the ability of splenic neutrophils from mice with liver metastases to suppress T-cell proliferation was reduced by blockade of IL-6R (Supp.Fig.4K). These data demonstrate that liver metastases induce changes in neutrophil state, with liver-derived IL-6 influencing this phenotype.

To obtain deeper insights into the difference in neutrophils depending on their origin and the presence or absence of metastases in different locations, we performed scRNA-seq. Neutrophils were isolated from the spleen and bone marrow in mice that were either tumour-free (naïve) or had metastases in liver or subcutaneous sites. Splenic neutrophils grouped into four clusters, with clusters 1 and 2 having higher expression of early neutrotime genes and clusters 3 and 4 having elevated expression of late neutrotime genes (Fig.4G, Supp.Fig.4L)^26^. Of note, clusters 1 and 2 expanded greatly when a liver metastasis was present, compared to mice with subcutaneous lesions and tumour-free mice. Further analysis indicated clusters 1 & 2 were similar to the G1, G2, and G3 neutrophil clusters reported by Xie *et al,* and were related to pre-neu, and immature classes described by Ervard and colleagues, while cluster 3 reflected a more mature G5a-c neutrophil phenotype (Supp.Fig.4M)^27,28^. Accordingly, neutrophils from spleens with liver metastases present had a G0-G3, immature phenotype (Supp.Fig.4N). Intriguingly, neutrophils in the spleen with a subcutaneous metastasis present expressed genes in G4 and G5a-c whereas naïve spleens expressed G5a genes, suggesting differences in more mature neutrophil phenotypes depending on the presence of cancer (Supp.Fig.4N). Moreover, despite the clustering being determined by genes independent of the cell cycle, cluster 1 alone had high expression of genes associated with S or G2 phases of the cell cycle, indicating that it reflects proliferating neutrophil progenitors (Fig.4H). In addition, G2M checkpoint and mitotic spindle pathways were upregulated in splenic neutrophils when a liver metastasis was present (Supp.Fig.4O), which is consistent with the EdU labelling (Fig.4A-D). Thus, the presence of liver metastases drives the increased production of neutrophils in the spleen with a distinctive early neutrotime G1-3 phenotype (Fig.4I, Supp.Fig.4N).

### Knockout of S100A9 enables the effective treatment of mice with liver metastases

To identify potential mediators of the distinct functionality of splenic-origin neutrophils when liver metastases are present, we further explored our single cell data, comparing differential gene expression splenic neutrophils of mice with or without melanoma liver metastases present. We observed numerous differentially expressed genes, including early neutrotime genes, G1-3 genes, genes associated with granules (*Ngp*, *Elane*, *Mpo*), *S100a8*, and *S100a9* (Fig.5A). In contrast, very few genes were differentially expressed in bone marrow neutrophils (Supp.Fig.5A), with similar neutrotime signatures between naive mice and mice bearing subcutaneous or liver metastases, indicating that liver metastases have a disproportionate effect on granulopoiesis in the spleen (Supp. Fig.5B). Flow cytometry further validated that the level of S100A9 was significantly higher in splenic neutrophils compared to bone marrow neutrophils (Fig.5B). Finally, in comparison to subcutaneous metastases, we observed higher gene and protein expression of S100A9 in liver metastases (Fig.5C and Supp.Fig.5C, gating strategy Supp.Fig.5D). We therefore examined the effect of *S100a9* knockout (KO), which also reduces S100A8 protein levels^29,30^, on the response of liver metastases to immunotherapy. We confirmed that S100A9 protein was lost from neutrophils in the KO mice (Supp.Fig. 5E&F). Loss of S100A9 did not reduce neutrophil numbers in the spleen or bone marrow (Supp.Fig.5G&H) and even increased neutrophil numbers in non-involved liver and liver metastases (Fig.5D&E).

**Figure 5.**
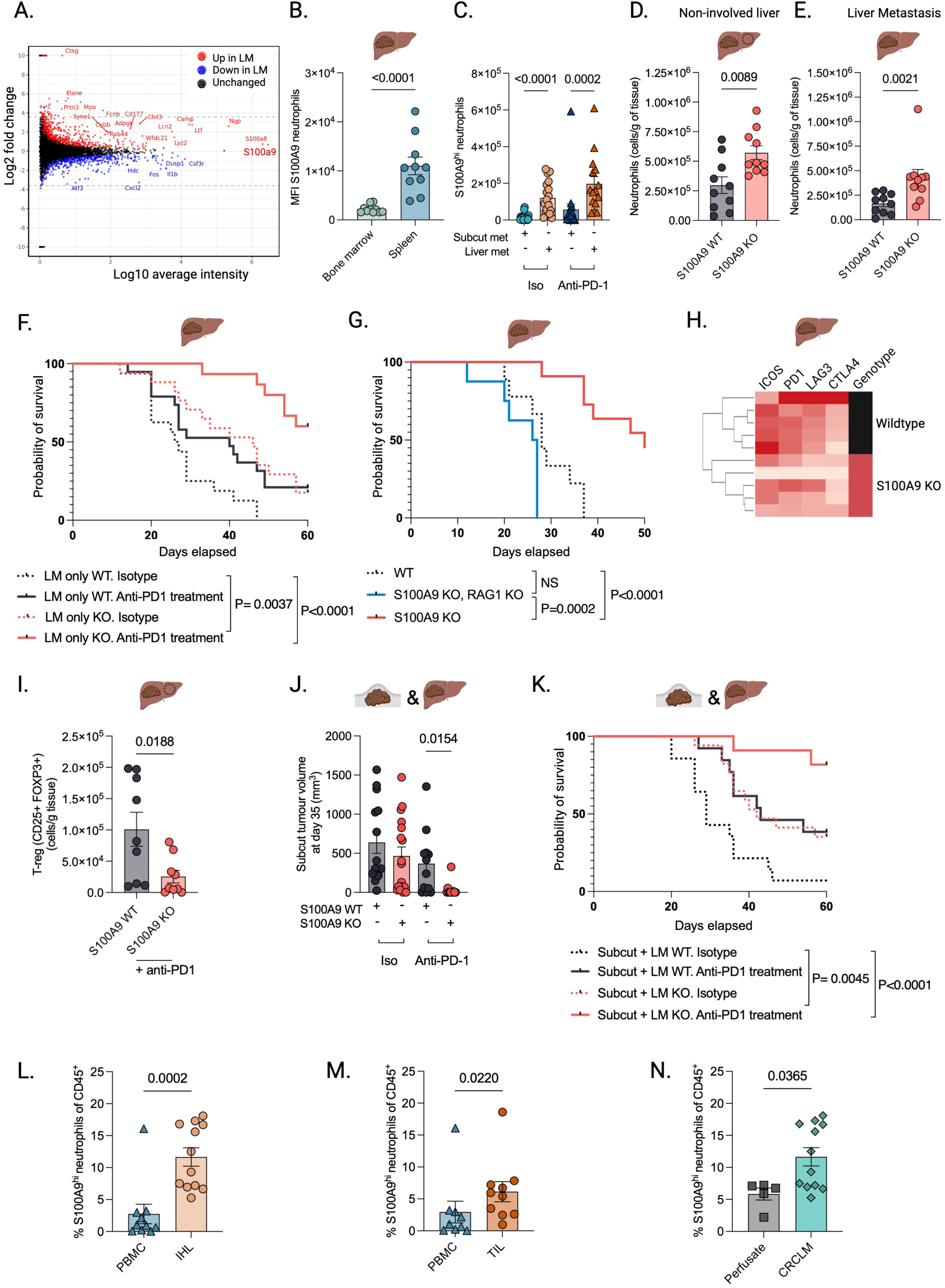
Knockout of S100A9 in combination with anti-PD-1 therapy results in improved survival of mice with subcutaneous and liver metastases. **A.** MA plot depicting differential gene expression of single cell sequenced splenic neutrophils in mice bearing liver metastases (LM) compared to naive liver (red=increased, blue=decreased, black=unchanged expression, n=5 biological replicates per group). **B.** Mean florescent intensity (MFI) of S100A9 in Ly6G^+^Ly6C^low^ neutrophils present in spleen vs. bone marrow of mice with liver metastases treated with isotype control (n=2 experiments). Mann-Whitney test used to test differences between groups. **C.** Absolute count of S100A9^hi^Ly6G^+^Ly6C^low^ neutrophils present in subcutaneous (subcut) and liver metastases treated with isotype control (iso) or anti-PD-1 (n=2 experiments). Kruskal-Wallis used to test differences between groups. **D&E.** Absolute count of Ly6G^+^Ly6C^low^ neutrophils present in **D.** Non-involved liver (with a LM present) and **E.** liver metastases in S100A9 knockout (KO) vs. wildtype (WT) mice with a liver metastasis present (n=2 experiments). Mann-Whitney test performed. F. Kaplan Meier survival estimate of wildtype (black) or S100A9 KO mice (orange) injected intrahepatically and treated with isotype control (iso, dashed line) or anti-PD-1 (continuous line). Differences in survival determined by log-rank test. G. Kaplan Meier survival estimate of RAG1 KO S100A9 KO (blue) vs. S100A9 KO (orange) vs. wildtype mice (black dotted line) with liver metastases. Differences in survival determined by log-rank test. H. Heatmap dendrogram clustering samples based on differential protein expression (mean florescence intensity using flow cytometry) of exhaustion markers (ICOS, PD1, LAG3, CTLA4) expressed by CD8^+^ T cells in liver metastases of S100A9 KO versus wildtype mice (n=5 biological replicates). I. Absolute count of CD4^+^CD25^+^FOXP3^+^ T_REG_ present in non-involved liver of S100A9 knockout (KO; orange) vs. wildtype (WT; black) mice with liver metastases present (n=2 experiments). Mann-Whitney test used to test differences between groups. J. Subcutaneous tumour volume (mm^3^) in wildtype (black) or S100A9 KO (orange) mice with liver metastases present. Unpaired T-test used to test differences between KO and WT anti-PD-1 treated groups. K. Kaplan Meier survival estimate of wildtype (black) or S100A9 KO mice (orange) with both liver and subcutaneous metastases present treated with isotype control (iso, dashed line) or anti-PD-1 (continuous line). Differences in survival determined by log-rank test. **L&M.** Frequency of human S100A9^hi^ low density neutrophils (HLA-DR^low^ CD33^+^ CD11b^+^ CD16^+^ ;as a proportion of live CD45^+^ leukocytes) in **L.** intrahepatic leukocytes (IHL) vs. peripheral blood (PBMC), **M.** tumour infiltrating lymphocytes (TIL) vs. PBMC of patients undergoing surgical resection for CRC liver metastases. Mann-Whitney test used to test differences between groups. **N.** Frequency of S100A9^hi^ neutrophils (HLA-DR^low^ CD33^+^ CD11b^+^ CD16^+^; as a proportion of live CD45+ leukocytes) in IHL of patients with CRC liver metastases compared to IHL obtained by perfusion of healthy donor livers prior to solid organ transplantation. Mann-Whitney test used to test differences between groups. All error bars represent mean ±S.E.M.

Next, we sought to understand how loss of S100A9 affected the growth and therapy response of the metastases. Mice lacking S100A9 with liver metastases showed decreased tumour growth (Supp.Fig.5I-J) and improved survival compared to age and sex matched co-housed wild-type mice (Fig.5F). Moreover, this survival was further boosted by administration of PD-1 antibodies (Fig.5F). Subcutaneous metastases in *S100a9* KO mice had slightly reduced survival compared to WT mice, but there was no difference in anti-PD-1 treated, as all responded (Supp.Fig.5K). To assess whether S100A9 high (S100A9^hi^) neutrophils affect adaptive immune responses to liver metastases, we established a *Rag1* KO, *S100a9* KO model. Liver metastases grew rapidly in these mice, resulting in significantly reduced survival compared to mice with *S100a9* KO only (Fig.5G). Depletion of CD8^+^ T cells confirmed that they were necessary for the improved survival of *S100a9* KO mice, thus suggesting that S100A9 enables neutrophils to suppress CD8^+^ T-cell function in liver metastases (Supp.Fig.5L). Consistent with this, CD8^+^ T cells in liver metastases growing in *S100a9* KO mice had reduced levels of the exhaustion markers PD1 and LAG3 (Fig.5H). A further observation in the *S100a9* KO animals was reduced numbers of T_REG_ compared to wildtype in the non-involved liver when liver metastases were present (Fig.5I). Further characterisation of these T_REG_ revealed that they largely consisted of a tissue resident (CXCR6^+^, CD69^+^) population, which had high CD44 expression suggesting a more immune suppressive phenotype^31^ (Supp.Fig.5M). These data indicate that *S100a9* is required for suppression of CD8+ T-cell functionality by liver metastases, and additionally suggest a role for T_REG_ in this process.

Liver metastases also prevent the response of distant lesions to immunotherapy, with subcutaneous metastases negatively impacted in our experimental model (Fig.1C), recapitulating observations in patients. We therefore evaluated the response of subcutaneous lesions to anti-PD-1 therapy when a liver metastasis was present. Strikingly, *S100a9* KO restored CPI efficacy, with greatly decreased subcutaneous tumour growth and significantly improved survival (Fig.5J&K, Supp.5N). Taken together, these data suggest S100A9 is crucial to both the local and systemic immunosuppression mediated by neutrophils when a liver metastasis is present

Finally, we assessed neutrophil S100A9 expression in human CRC liver metastases. We observed higher S100A9 expression in neutrophils in both IHL and TIL compared to the periphery (Fig.5.L&M, gating strategy Supp.Fig.5P). Critically, we observed higher numbers of S100A9^hi^ neutrophils in CRC liver metastases compared to perfusates taken from healthy donor transplant livers, suggesting the presence of liver metastasis is associated with neutrophils with high levels of S100A9 (Fig.5N).

## Discussion

Liver metastases present a particular challenge for immunotherapy, with only half of patients with melanoma liver metastases in our cohort surviving for 12 months. Through an experimental model that recapitulates the CPI refractory nature of liver metastases, we identified the production of S100A9^hi^, early ‘neutrotime’, G0-3 neutrophils in the spleen as a critical factor determining why liver metastases reduce the efficacy of CPI. Within the liver microenvironment, these neutrophils suppress T-cell function and their presence is associated with increased T-cell exhaustion, resulting in impaired anti-tumour activity and resistance to CPI. In addition, once these neutrophils leave the spleen, they may transit to other sites of metastases leading to systemic suppression of anti-tumour responses (Fig. 6). These immature S100A9^hi^ neutrophils directly suppress the proliferation and activation of CD8^+^ T cells. Based on our imaging data, we propose that these interactions occur at the margins of liver metastases and in the liver parenchyma. Furthermore, *S100a9* KO reduced numbers of tissue resident T_REGS_, suggesting that S100A9^hi^ neutrophils could also influence systemic immune responses to CPI through modulating interactions between CD8^+^ Tcells and T_REG_. Recent characterisation of tissue resident T_REG_ has shown that contrary to prior assumptions, they have only transient residency and re-circulate throughout the rest of the organism^32^. Thus, in addition to systemic dissemination of splenic neutrophils, our data support the observations that transit of T_REG_ from the liver to other sites might contribute to systemic immune suppression^12^. Interestingly, we did not see evidence within our models or patient data of macrophage-mediated CD8^+^ T-cell elimination previously reported in liver metastases^20^.

**Figure 6.**
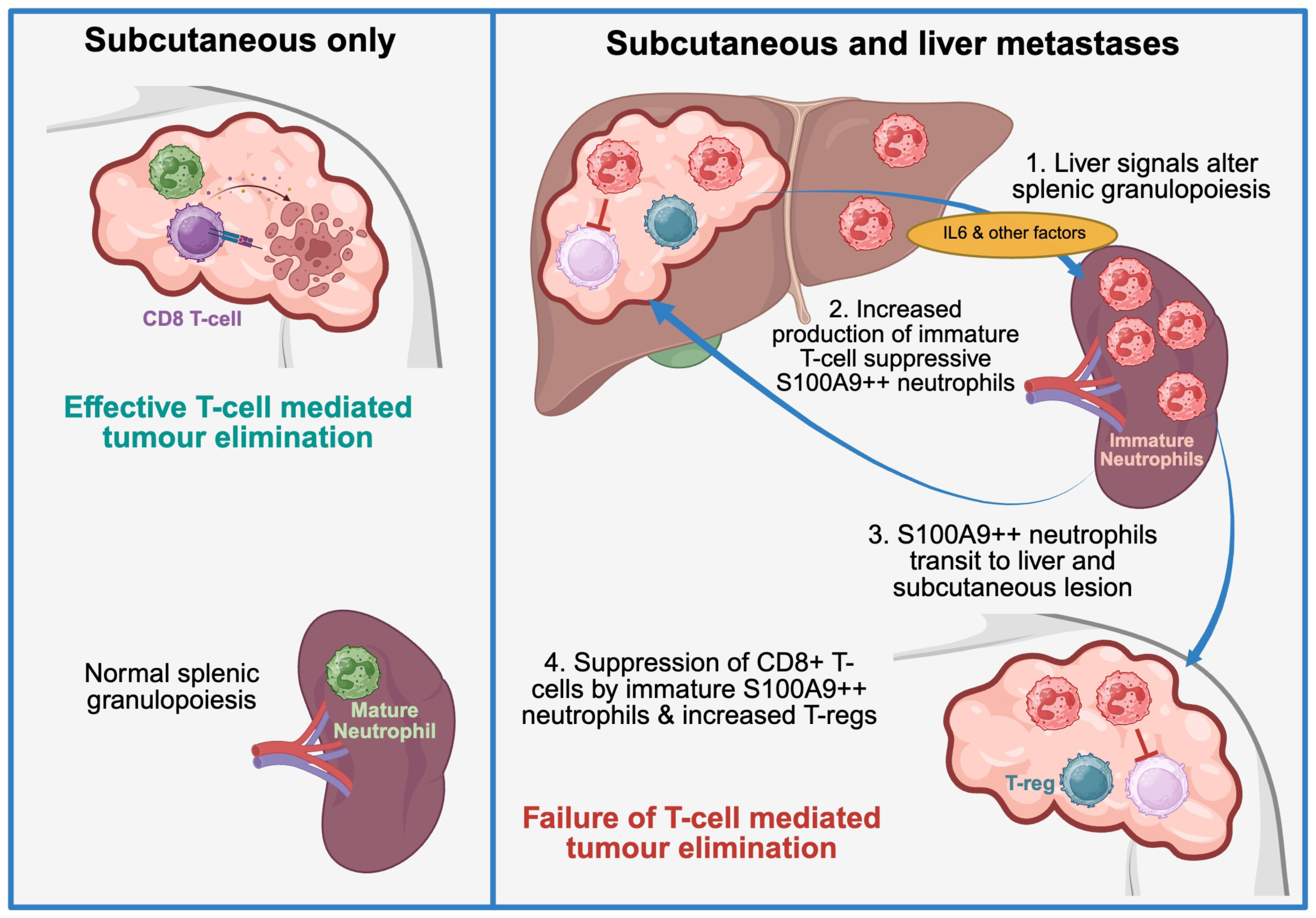
**Overview of liver-induced splenic granulopoiesis resulting in production of immune-suppressive neutrophils** Green neutrophils= tumour inhibitory, red neutrophils=tumour promoting

There is increasing recognition that neutrophil populations/states can be very heterogenous, with certain subsets having an anti-tumour role whilst others being pro-tumorigenic^33,34^. In addition, there is increasing evidence that splenic neutrophils may have different developmental maturation rates and transcriptional programmes compared to bone marrow neutrophils^35^. We now show that the origin of neutrophils and the drivers of their proliferation are important to their immune suppressive state. Liver metastases predominantly caused changes in both proliferation and phenotype of splenic, rather than bone marrow-derived, neutrophils, with early neutrotime, G1-3 markers and S100A9 expression being particularly high in this context. Subcutaneous metastases did not lead to changes in splenic granulopoiesis and were infiltrated with neutrophils with higher IFNγ-linked gene expression. This observation is consistent with more mature, IFNγ responsive, Ly6E high neutrophils^26^ being associated with improved outcomes to CPI^36^. These observations sit in the wider context of studies suggesting that the presence of cancer, particularly liver metastases, results in increased numbers of immature neutrophils^28,37^. Neutrophils also play an important role in non-alcoholic steatohepatitis driven hepatocellular carcinoma, although the relationship between neutrophil maturity and function may be different in this context^38^. The splenic granulopoiesis triggered by liver metastases in likely to be distinct from emergency granulopoiesis in response to infection, which previous work has shown matures neutrophil populations towards a G4 and G5b phenotype particularly in the peripheral blood and spleen^27^. Importantly, it raises the possibility that driving neutrophils towards a more mature, IFNγ responsive, anti-tumour state would be desirable. The validity of this approach has been demonstrated in mice through reduced 4T1 tumour growth following the combination of anti-PD-1 with adoptive transfer of Ly6E high neutrophils^36^, although it remains to be determined if this approach would be viable in patients. Thus, better understanding of how neutrophil development could be driven along maturation trajectories will be critical to generation of new targets.

Work by Evrard *et al* showed that pre-neu in the bone marrow have lower levels of *S100a8* than more mature neutrophils^28^. Intriguingly, expression of *S100a8/9* was high in our splenic neutrophils with liver metastases present despite also being associated with a more immature, G1-3 pre-neu/im-neu phenotype. This suggests that the timing of gene expression events during neutrophil maturation when a liver metastasis is present might be disrupted in the spleen, or that neutrophil maturation may be wired differently in the spleen and bone marrow. Supporting the latter hypothesis, we did not observe high levels of *S100a9* in bone-marrow derived neutrophils when liver metastases were present. S100A8 and S100A9 expression on neutrophils has been associated with tumour progression in many cancers and serum levels have been associated with poor outcomes to anti-PD-1 therapy in patients with melanoma, suggesting this could be an important target to improve CPI response ^39–41^.

Our analyses raise several pertinent questions. The molecular means of communication between the liver and spleen remains to be fully elucidated. Although anti-IL-6R reduced splenic neutrophil numbers, this was not to the same level as naïve mice. We speculate that this may reflect redundancy in signalling between the liver and spleen, with communication between the organs having evolved to have no single point of failure. It will be interesting to more fully map out how liver metastases modulate granulopoiesis in the spleen. In addition, it will be important to determine how to achieve similar effects to *S100a9* knock-out in patients. The functions of S100A9 are multifaceted and include migration, activation and NET production^42,43^. Further, they can be released and act as damage-associated molecular pattern (DAMP) mediators, resulting in an inflammatory response^44,45^. Therefore, it is unlikely that targeting one specific downstream pathway will be sufficient to mediate an overall effect. Recent advances in proteolysis-targeting chimera (PROTAC) protein degraders may offer a route to clinical translation^30^. Importantly, knockout of *S100a9* does not impair the viability or health of mice and previous studies have shown that important antimicrobial neutrophil functions such as phagocytosis, superoxide burst, and apoptosis are unaffected by *S100a9* depletion, suggesting that there may be a therapeutic window in which to target S100A9^hi^ neutrophils in patients ^46^.

In summary, we have shown that the presence of liver metastasis has important effects on the host immune system, driving splenic granulopoiesis which results in production of particularly immune suppressive neutrophils. Our data further support the rationale that precision immunotherapy should take into consideration the organ microenvironment of the metastasis as well as tumour characteristics. Crosstalk between the tumour and the host can result in tolerising effects beyond the local tumour microenvironment. Targeting mediators of both local and systemic tolerance such as S100A9^hi^ neutrophils may enable better outcomes to immune therapy.

## Methods

### Cell culture

All cell lines were provided by the Cell Services Unit of The Francis Crick Institute, where they were authenticated using Short-Tandom Repeat (STR) profiling and species-identification tests, and confirmed to be mycoplasma-free. 4434 mouse melanoma (donated by Marais laboratory), CT26 (colorectal) or 4T1 (breast) cells were cultured in DMEM (ThermoFisher, #41966052) containing 10% fetal bovine serum (Gibco, #10270-106), 1% penicillin/streptomycin (Invitrogen, #15140122) and kept at 37°C and 5% CO_2_.

### Mice

Female 6-12 week-old, C57BL/6 mice were used for all experiments or wildtype controls. *S100a9* KO or *S100a9* KO, *RAG1* KO animals were female, age matched and co-housed with wildtype controls. For all immunotyping experiments, mice were culled at day 14. Subcutaneous tumours were measured using calipers and liver ultrasound was performed using a Vevo 3100 ultrasound (Fujifilm visualsonics). In treatment experiments, anti-PD-1 clone RMP1-14 200mg/kg or anti-Gr-1 400mg/kg or anti-IL-6R, clone 15A7 with corresponding isotype controls (rat IgG2a or rat IgG2b respectively) were given from day 3 twice weekly via intra peritoneal injection for up to 6 doses (all BioXcell). Further neutrophil depletion experiments used 50mg/kg anti-Ly6G, clone 1A8 followed 2 hours later by 50mg/kg anti-rat anti-rat Kappa Immunoglobulin Light Chain, both given via intraperitoneal injection, three times a week for up to 3 weeks. In addition, CD8 depletion experiments were performed using 500μg per mouse anti-CD8α (clone 2.43) or rat IgG2b isotype control (BioXcell) from day 3, twice weekly via intra peritoneal injection for up to 6 doses. Mice were euthanised at day 14 for downstream immunophenotying experiments. Experiments were approved by the Animal Welfare and Ethical Review Body of the Francis Crick Institute and authorized by the U.K. Home Office (PPL: PP0736231).

### Mouse cell isolation

Livers, spleens, tumours, blood and bone marrow were collected from mice in phosphate-buffered saline (PBS), and lymphocytes were isolated by gentle mechanical disruption through a 70-μm cell strainer (Greiner). Intrahepatic cells were isolated using 30% Percoll (GE Healthcare) density gradient centrifugation to remove dead cells and hepatocytes. Red cell lysis was then performed by exposure to red cell lysis buffer (Miltenyi Biotec) for 3 minutes prior to PBS wash and centrifugation.

### Human cell isolation

IHLs were isolated from resected/explanted liver material as previously described^47^. In brief, tissue was cut up and incubated at 37°C for 30 min in HBSS^+/+^ (Life Technologies; Thermo Fisher Scientific) containing collagenase IV (Thermo Fisher Scientific) and DNase I (Roche; Sigma-Aldrich; Merck), before mechanical disruption using the GentleMACS (Miltenyi Biotech) dissociator, with debris removed by filtration through a 70 µm filter (Greiner). Parenchymal cells were removed by centrifugation on a 30% Percoll gradient (GE Healthcare; VWR) followed by further leukocyte isolation by density centrifugation using a Pancoll gradient (PAN Biotech). IHLs were isolated from perfusion liquid (perfusates) or small core-biopsy samples as previously described^19^. In brief, the perfusate fluid was first concentrated by centrifugation. Concentrated cells were then resuspended in RPMI-1640 (Life Technologies; Thermo Fisher Scientific) and isolated by density centrifugation on a Pancoll gradient.

### Immunophenotyping

Multiparametric flow cytometry was used for phenotypic and functional analysis of mouse/human lymphocytes/myeloid cells. Cells were stained with a blue fixable Live/Dead dye (Invitrogen; Thermo Fisher Scientific) for the exclusion of dead cells before incubation with saturating concentrations of surface monoclonal antibodies (mAbs) diluted in 50% Brilliant violet buffer (BD Biosciences) and 50% PBS for 30 min at 4°C (Full details of the monoclonal antibodies, including dilutions and catalogue numbers, are provided in Key Resources table). Cells were fixed and permeabilized for further functional assessment with Cytofix/Cytoperm (BD Biosciences) or the Foxp3 buffer kit (BD Bioscience) according to the manufacturer’s instructions. For intracellular cytokine staining, samples were then incubated with saturated concentrations of intracellular mAbs for 30 min at 4°C diluted in 0.1% saponin (Sigma-Aldrich) or 1× PBS for 30 min at 4°C. All samples were acquired in 1x PBS on BD Fortessa X20 flow cytometer (BD Biosciences) running DIVA (v.8.0.1). Data were analysed using FlowJo v10.10.0 (Tree Star). Doublets and dead cells were excluded from the analysis.

### Proliferation analysis with EdU

The Click-iT™ EdU Alexa Fluor™ 647 Flow Cytometry Assay Kit (Invitrogen #C10419) was used to perform the assay. Mice were injected with 4434 cells in the locations described and at day 14 of tumour growth, mice were injected intraperitoneally with EdU (25mg/kg) 1 hour prior to being culled. Cells were stained as per manufacturers’ instructions with the addition of neutrophil surface mAbs (for gating see Supp.4A).

### Single cell analysis

Mice were anaesthetised and the inferior vena cava was cannulated, and collagenase perfused into the liver using a pump at a flow rate of 5ml/min. Tumours were dissociated using 1/66 TH Liberase, 1/66 TM Liberase and 1/400 DNAse solution for 30 mins at 37°C. Cells were then isolated as above. For experiments sequencing all cells, these were first sorted for live cells. The concentration and viability of the single cell suspension was measured using acridine orange and propidium iodide and trypan blue on the Luna-FX7 Automatic Cell Counter. Approximately (5000-20,000) cells were loaded on Chromium Chip and partitioned in nanolitre scale droplets using the Chromium Controller and Chromium Next GEM Single Cell Reagents. Within each droplet the cells were lysed, and the RNA was reverse transcribed. All the resulting cDNA within a droplet shared the same cell barcode. Illumina compatible libraries were generated from the cDNA using Chromium Next GEM Single Cell library reagents in accordance with the manufacturer’s instructions. Final libraries were checked using the Agilent TapeStation and sequenced using the Illumina NovaSeq 6000. Sequencing read configuration: 28-10-10-90.

For experiments examining neutrophils only, these were first surface stained with mAbs then fixed using the 10x Flex fixation protocol according to manufacturers’ instructions. Cells were sorted (for gating strategy see Supp.5P) Samples were fixed in a 4% formaldehyde fixative solution using Chromium Next GEM Single Cell Fixed RNA Sample Preparation Kit (10x Genomics). Gene expression was measured using barcoded probe pairs designed to hybridize to mRNA specifically. Using a microfluidic chip, the fixed and probe-hybridized single cell suspensions were partitioned into nanolitre-scale Gel Beads-in-emulsion (GEMs). A pool of ∼737,000 10x GEM Barcodes was sampled separately to index the contents of each partition. Inside the GEMs, probes were ligated and the 10x GEM Barcode was added, and all ligated probes within a GEM share a common 10x GEM Barcode. Barcoded and ligated probes were then pre-amplified in bulk, after which gene expression libraries were generated according to manufacturer’s instructions and sequenced (NovaSeq 6000. Sequencing read configuration: 28-10-10-90).

Raw single-cell sequencing data from individual samples were processed using the 10x CellRanger pipeline (10x Genomics) and analysed with the Seurat R-package (version 5.0.0)^48^, in R version 4.3.1. Cells were filtered to exclude cells with a mitochondrial gene percentages of higher than 12.5% and cells with less than 200 RNA features per cell. To create the UMAP plots, the first 30 PCA dimensions were included and a partial cell-cycle regression to equalize all G2M and S-phase differences was performed. Single-cell differential gene expression analyses were conducted with the glmGamPoi R-package (version 1.12.2)^49^.

### Neutrophil and T cell co-culture experiments

Mouse and spleen neutrophils were isolated as above then incubated in FcR blocking reagent (Miltenyi) for 10 minutes at 4°C. PE-conjugated anti-Ly6G antibody was added at 1:100 and incubated on ice for 30min. After washing with MACS buffer, cells were incubated with anti-PE-conjugated microbeads (Miltenyi Biotec) for 15 min at 4°C. Positive cell selection was performed using magnetic separation columns (Miltenyi Biotec). T cells were isolated from different spleens using EasySep™ Mouse T Cell Isolation Kit as per manufacturer’s protocol. T cells were stimulated using Dynabeads™ Mouse T-Activator CD3/CD28 for T Cell Expansion and Activation using manufacturer’s protocol. T cells and neutrophils were co-cultured 1:1 overnight for short term stimulations or up to 7 days for assessment of proliferation before antibody staining and flow cytometry.

### Imaging mass cytometry (IMC)

IMC panel development and staining was performed as described in Giangreco et al^35^. In brief, custom antibodies were validated with immunofluorescence staining and conjugated to heavy metal tags using Maxpar-X8 antibody labelling kits (Standard Biotools) according to manufacturer’s protocol. For tissue microarray staining, FFPE sections were dewaxed, antigen retrieval was performed using Tris-EDTA Ph9 buffer and slides were stained overnight at 4’C with metal-conjugated antibodies. Slides were stained with Iridium and Ruthenium counterstains and left to dry before imaging on a Hyperion CyTOF imaging system. Representative images were obtained using ImageJ2 2.16.0 (Fiji).

Cell segmentation and phenotyping was performed using the TRACERx-PHLEX pipeline^50^ as described in Giangreco et al^51^. In brief, nuclei were segmented using Iridium channels and expanded by 1µm to create cell objects with the deep-imcyto module. Cell objects were then phenotyped according to expression of major lineage markers using the TYPEx module. 10-cell neighbourhoods were identified using the Spatial-PHLEX module and k-means unsupervised clustering of neighbourhoods was performed using the ClusterR package in R (RStudio version 4.3.1). Further downstream data analysis was performed in R (RStudio version 4.3.1).

### Patient samples and data

Blood and liver samples were obtained through the Tissue Access for Patient Benefit Initiative (TAPb) at The Royal Free Hospital (approved by the UCL-Royal Free Hospital BioBank Ethical Review Committee references: 11/WA/0077, 16/WA/0289 and 21/WA/0388); healthy donor blood samples were obtained through REC 11//LO/0421. Clinical data were obtained under UK Computer-Aided Theragnostics (ukCAT) REC 21/NW/0347 from patients attending The Christie Hospital treated first line for advanced disease with anti-PD-1 or anti-PD-1 plus CTLA-4 therapy. Median OS with 95% confidence intervals (CIs) by liver metastasis, lymphocyte, NLR or neutrophil status was estimated using the Kaplan–Meier method. Hazard ratios (HRs) with 95% CIs were estimated using Cox proportional hazards models, adjusting for treatment, cancer type, age, sex, performance status, baseline LDH, and baseline neutrophil-to-lymphocyte ratio. Subgroup analyses were performed by cancer type, with melanoma models additionally adjusted for BRAF mutation status. All survival analyses were conducted using R (version 4.4.0).

## Key Resources table

| REAGENT or RESOURCE | SOURCE | IDENTIFIER |
| --- | --- | --- |
| **Antibodies** |  |  |
| BD Horizon™ BUV395 anti-human HLA-DR Antibody (clone L243) | BD Biosciences | Cat#568648; RRID: AB_3684432 |
| BD Horizon™ BUV805 anti-Human CD45 Antibody (clone HI30) | BD Biosciences | Cat#612891; RRID: AB_2870179 |
| Brilliant Violet 510™ anti-human CD14 Antibody (clone M5E2) | Biolegend | Cat#301842; RRID: AB_2561946 |
| FITC anti-human MRP-14 (S100A9) FITC Antibody (clone MRP 1H9) | Biolegend | Cat#350704; RRID: AB_10662250 |
| eBioscience™ PE-Cyanine7 anti-human CD11b Antibody (clone ICRF44) | Thermo Fisher Scientific | Cat#25-0118-42; RRID: AB_1582272 |
| BD Pharmingen™ Alexa Fluor 700® anti-human CD33 Antibody (clone WM53) | BD Biosciences | Cat#561160; RRID: AB_10561683 |
| BD Pharmingen™ APC-Cy™7 anti-human CD16 Antibody (clone 3G8) | BD Biosciences | Cat#557758 RRID: AB_396864 |
| LIVE/DEAD Fixable Blue Dead Cell Stain Kit | Thermo Fisher Scientific | Cat#L34961 |
| PerCP/Cyanine5.5 anti-mouse/human CD44 Antibody (clone IM7) | Biolegend | Cat#103032 RRID: AB_2076204 |
| Brilliant Violet 785™ anti-mouse CD4 Antibody (clone RM4-5) | Biolegend | Cat#100552 RRID: AB_2563053 |
| Brilliant Violet 711™ anti-mouse PD-1 Antibody (clone 29F.1A12) | Biolegend | Cat#135231 RRID: AB_2566158 |
| Brilliant Violet 605™ anti-mouse CD69 Antibody (clone H1.2F3) | Biolegend | Cat#104530 RRID: AB_2563062 |
| Brilliant Violet 510™ anti-mouse CD25 Antibody (clone PC61) | Biolegend | Cat#102041 RRID: AB_2562269 |
| Brilliant Violet 421™ anti-mouse CD186 (CXCR6) Antibody (clone SA051D1) | Biolegend | Cat#151109 RRID: AB_2616760 |
| BD Horizon™ BUV805 anti-mouse CD8a Antibody (clone 53-6.7) | BD Biosciences | Cat#612898 RRID: AB_2870186 |
| BD Horizon™ BUV395 anti-mouse CD3e Antibody (clone 145-2C11) | BD Biosciences | Cat#563565 RRID: AB_2738278 |
| APC/Cyanine7 anti-mouse CD45 Antibody (clone 30-F11) | Biolegend | Cat#103116 RRID: AB_312981 |
| Alexa Fluor® 700 anti-mouse CD62L Antibody (clone MEL-14) | Biolegend | Cat#104426 RRID: AB_493719 |
| Alexa Fluor® 647 anti-mouse FOXP3 Antibody (clone MF-14) | Biolegend | Cat#126408 RRID: AB_1089115 |
| PE/Cyanine7 anti-human/mouse/rat CD278 (ICOS) Antibody (clone C398.4A) | Biolegend | Cat#313520 RRID: AB_10643411 |
| PE/Dazzle™ 594 anti-mouse CD223 (LAG-3) Antibody (clone C9B7W) | Biolegend | Cat#125224 RRID: AB_2572082 |
| PE anti-mouse CTLA4/CD152 Antibody (clone UC10-4B9) | Biolegend | Cat#106306 RRID: AB_313255 |
| Brilliant Violet 605™ anti-mouse CD274 (B7-H1, PD-L1) Antibody (clone 10F.9G2) | Biolegend | Cat#124321 RRID: AB_2563635 |
| Brilliant Violet 510™ anti-mouse Ly6C Antibody (clone HK1.4) | Biolegend | Cat#128033 RRID: AB_2562351 |
| PerCP/Cyanine5.5 anti-mouse CD3 Antibody (clone 17A2) | Biolegend | Cat#100218 RRID: AB_1595492 |
| BD Horizon™ BUV805 anti-mouse CD45 Antibody (clone 30-F11) | BD Biosciences | Cat#748370 RRID: AB_2872789 |
| BD Horizon™ BUV395 anti-mouse NK1.1 Antibody (clone PK136) | BD Biosciences | Cat#564144 RRID: AB_2738618 |
| APC/Cyanine7 anti-mouse Ly6G Antibody (clone 1A8) | Biolegend | Cat#127624 RRID: AB_10640819 |
| eBioscience™ Alexa Fluor™ 700 CD11b Monoclonal Antibody (clone M1/70) | Thermo Fisher Scientific | Cat#56-0112-82 RRID: AB_657585 |
| S100A9 (D3U8M) Rabbit mAb (Rodent Specific) PE Conjugate | Cell Signalling Technology | Cat#93941 RRID: AB_2800218 |
| PE anti-mouse CD117 (c-kit) [2B8] | Biolegend | Cat#105807 RRID: AB_313216 |
| InVivoPlus anti-mouse Ly6G/Ly6C (Gr-1), RB6-8C5 | BioXcell | Cat#BP0075 |
| InVivoPlus rat IgG2b Isotype control | BioXcell | Cat#BP0090 |
| InVivoPlus anti-mouse PD-1 (CD279), clone RMP1-14 | BioXcell | Cat#BP0146 |
| InVivoPlus rat IgG2a isotype control, anti-trinitrophenol | BioXcell | Cat#BP0089 |
| InVivoPlus rat, anti-Mouse IL-6R, clone 15A7 | BioXcell | Cat#BE0047 |
| InVivoMAb anti-mouse Ly6G, clone 1A8 | BioXcell | Cat#BE0075-1 |
| InVivoMab anti-rat Kappa Immunoglobulin Light Chain | BioXcell | Cat#BE0122 |
| **Biological samples** |  |  |
| Human colorectal cancer and liver | TapB | N/A |
| **Chemicals, peptides, and recombinant proteins** |  |  |
| HBSS<sup>++</sup> | Thermo Fisher Scientific | Cat#24020-091 |
| DNase I | Sigma-Aldrich | Cat#11 284 932 001 |
| Collagenase Type IV | Thermo Fisher Scientific | Cat#17104019 |
| Percoll | VWR | Cat#17-0891-02 |
| RPMI | Thermo Fisher Scientific | Cat#21875-034 |
| DMEM | Thermo Fisher Scientific | Cat#11965092 |
| Pancoll human, Density 1.077 g/ml | Pan Bio Tech | Cat#P04-60500 |
| FCR blocking reagent | Miltenyi Biotec | Cat#130-059-901 |
| Brilliant Stain Buffer | BD Biosciences | Cat#566349 |
| Cytofix/Cytoperm Fixation and Permeabilisation solution | BD Biosciences | Cat#554722 |
| CountBright™ Absolute Counting Beads | Thermofisher | Cat#C36950 |
| Red Blood Cell Lysis Solution | Miltenyi Biotec | Cat#130-094-183 |
| Liberase™ TM | Sigma Aldrich | Cat#5401119001 |
| Liberase™ TH | Sigma Aldrich | Cat#5401135001 |
| PE-conjugated microbeads | Miltenyi Biotec | Cat# 130-048-801 |
| **Critical commercial assays** |  |  |
| EasySep™ Mouse T Cell Isolation Kit | STEMcell technologies | Cat#19851 |
| Dynabeads™ Mouse T-Activator CD3/CD28 | Thermofisher | Cat#11452D |
| Click-iT™ EdU Alexa Fluor™ 647 Flow Cytometry Assay Kit | Invitrogen | Cat#C10419 |
| Chromium Next GEM Single Cell Fixed RNA Sample Preparation Kit | 10x | Cat#PN-1000414 |
| Chromium Next GEM Single Cell 3' GEM, Library & Gel Bead Kit v3.1 | 10x | Cat#PN-1000121 |
| Fixed RNA Feature Barcode Kit 16 rxns | 10x | Cat#PN-1000419 |
| **Deposited data** |  |  |
| Raw and analysed data | This paper | Data will be provided upon reviewers request and will be uploaded to the GEO repository |
| **Experimental models: Cell lines** |  |  |
| 4434 mouse melanoma | Francis Crick cell services | N/A |
| CT26 mouse colorectal cancer | Francis Crick cell services | N/A |
| 4T1 mouse breast cancer | Francis Crick cell services | N/A |
| **Experimental models: Organisms/strains** |  |  |
| MRP-14 (S100a9 -/-) mouse | Manitz et al 2003 | N/A |
| MRP-14 (S100a9 -/-), Rag-1-/- mouse | This paper | N/A |
| C57BL/6Jax | Francis Crick Institute | N/A |
| **Software and algorithms** |  |  |
| Flowjo (version 10.10.0) | Becton, Dickinson and Company | https://www.flowjo.com/flowjo/overview |
| R (version 4.4.0 and 4.3.1) | R Core Team<sup>52</sup> | https://www.R-project.org/ |
| ImageJ2 (version 2.16.0) | Schneider et al<sup>53</sup> | https://imagej.nih.gov/ij/ |
| TRACERx-PHLEX | Magness et al<sup>50</sup> | https://www.nature.com/articles/s41467-024-48870-5 |
| **Other** |  |  |

## Supplementary Figures

**Supplementary Figure 1.**
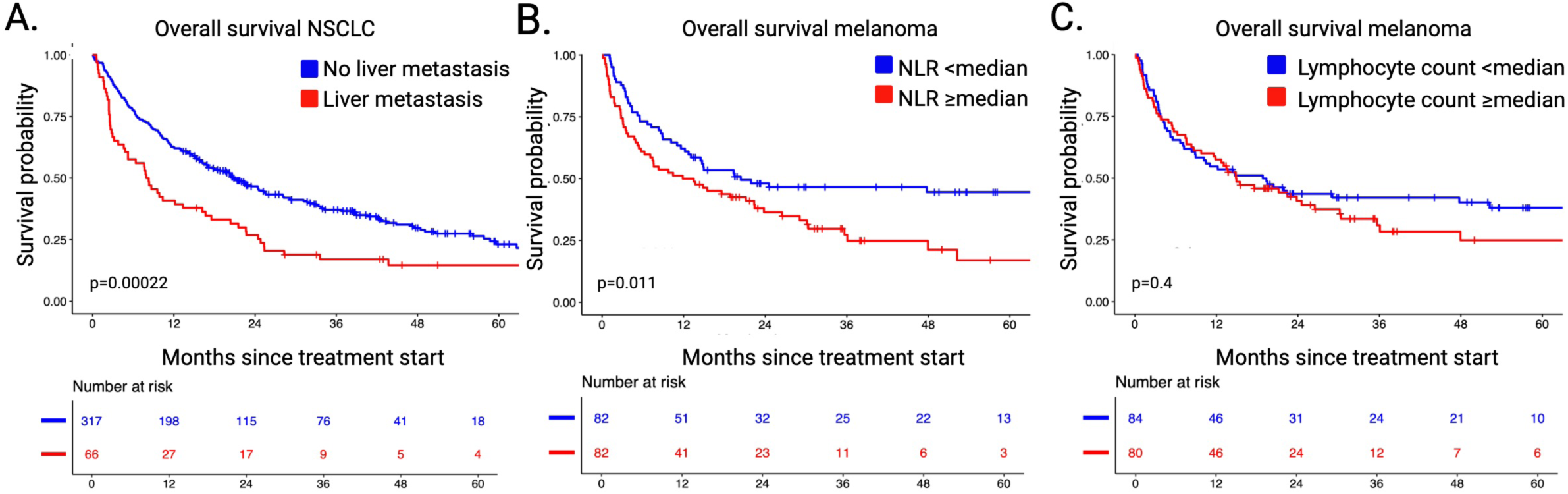
A. Kaplan Meier survival estimate of patients with non-small cell lung cancer (NSLC) metastasis (n=66, red) vs. without (n=317, blue) treated with an anti-PD-1 containing regimen were followed for a median of 48 months. Differences in survival determined by log-rank test. **B&C**. Kaplan Meier estimate of overall survival in patients with melanoma liver metastasis (n=164) treated with an anti-PD-1 containing regimen stratified by **B.** neutrophil lymphocyte ratio (NLR) and **C.** lymphocyte count of the cohort ≥median (red) vs. <median (blue). Differences in survival determined by log-rank test.

**Supplementary Figure 2.**
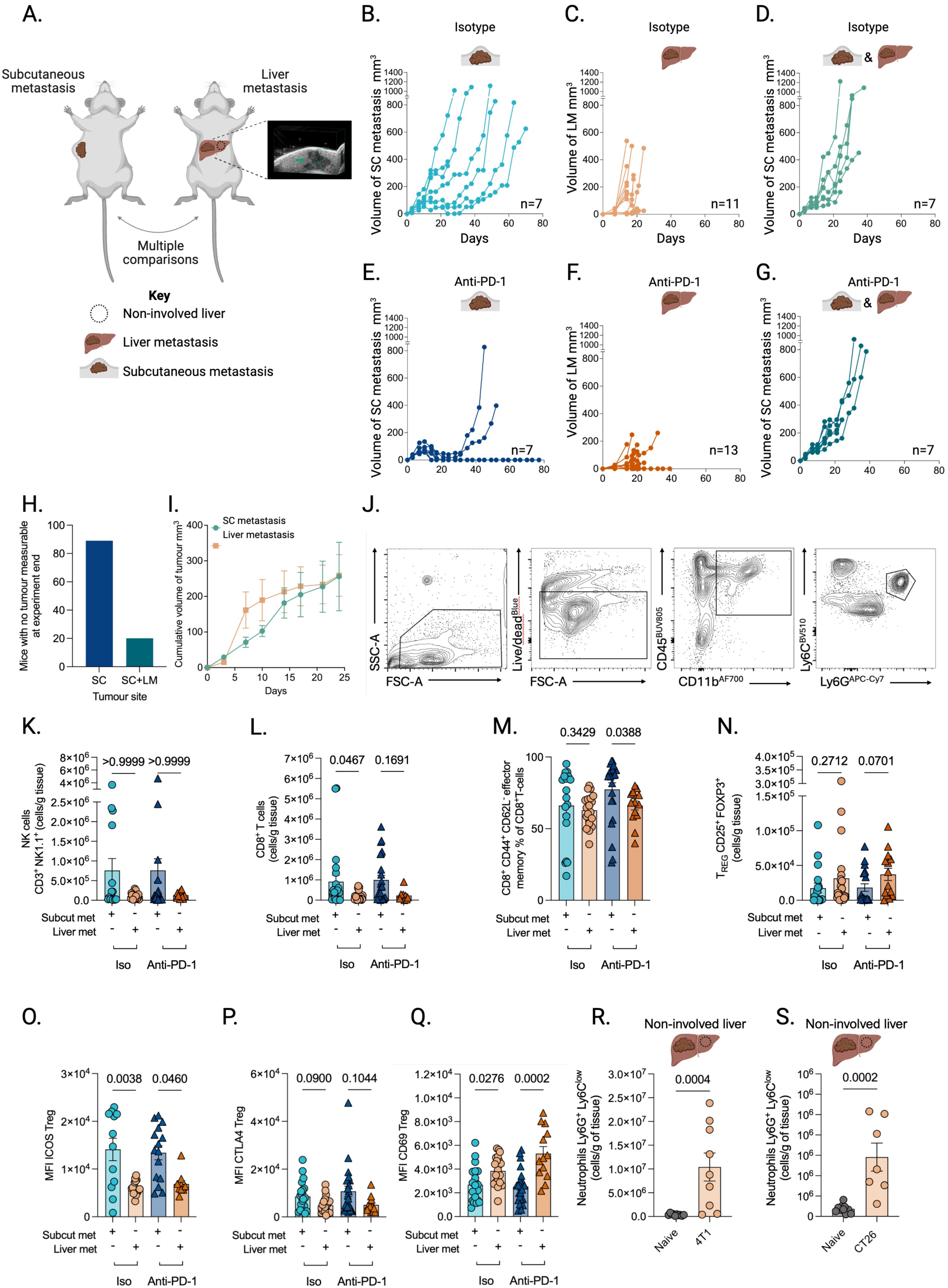
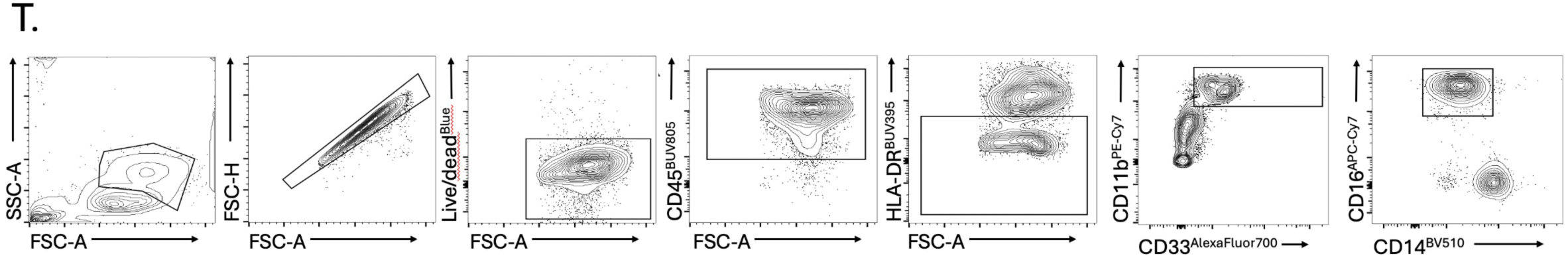
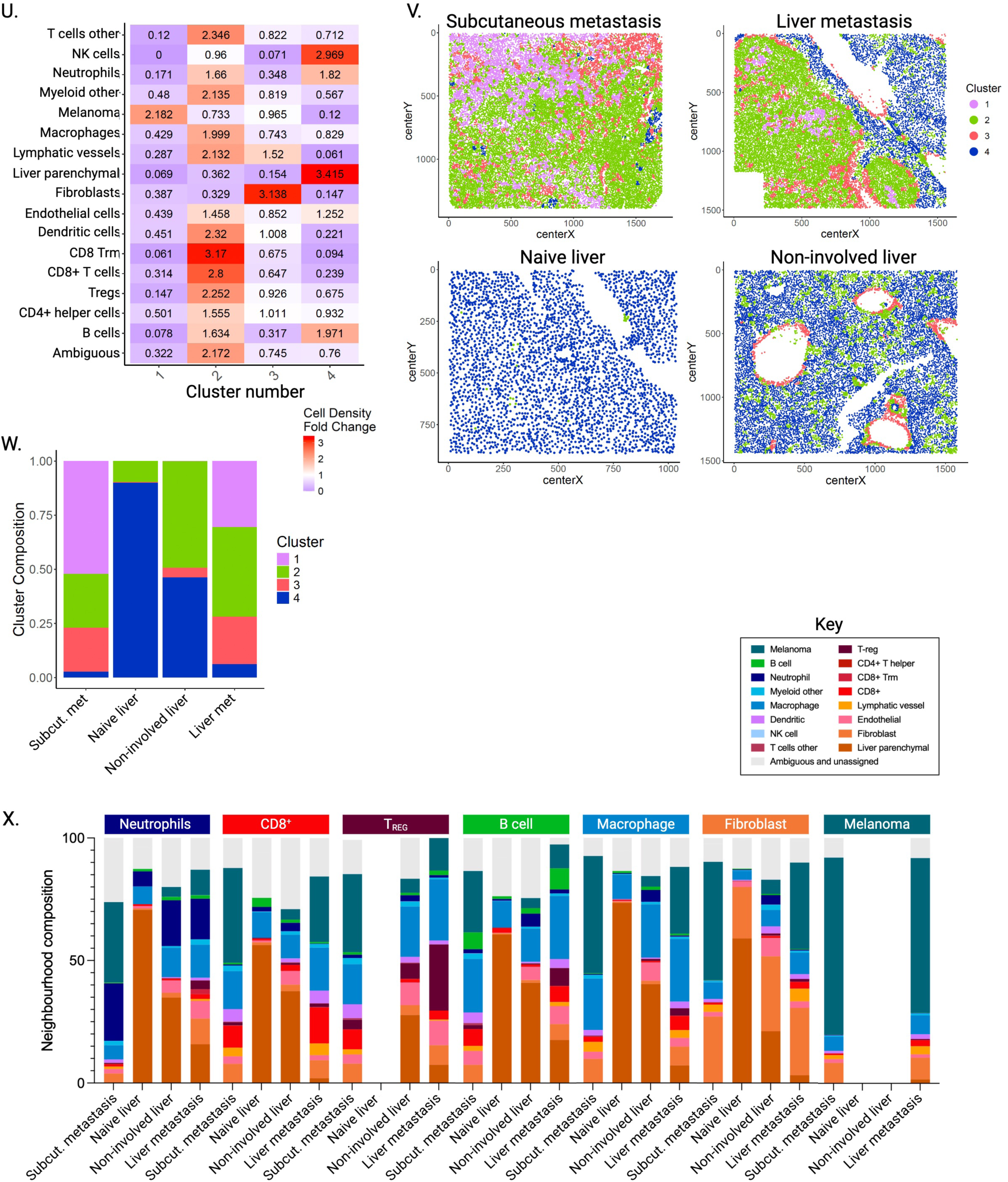
A. Experimental set-up. Immunocompetent C57/Bl6 mice were injected with 4434 *BRAF* mutant mouse melanoma cells either subcutaneously or orthotopically into the liver using ultrasound resulting in discrete tumours. **B-G.** Growth of **B.** subcutaneous metastasis measured using calipers **C.** liver metastasis measured using ultrasound **D.** both subcutaneous and liver metastases treated with isotype control. Growth of **E.** subcutaneous metastasis **F.** liver metastasis **G.** both subcutaneous and liver metastases treated with anti PD-1. Of note subcutaneous growth endpoint affected if mice die from LM. **H.** Frequency of complete response in the subcutaneous site with vs. without liver metastases present. **I.** Cumulative volume of tumours in mice with subcutaneous metastasis measured using calipers and liver metastasis measured using ultrasound (n=7 biological replicates per group). **J.** Sequential gating strategy to define mouse neutrophils (CD45^+^ CD11b^+^ Ly6G^+^ Ly6C^low^) from freshly isolated lymphocytes. **K&L.** Absolute count of **K.** CD3^+^ NK1.1^+^ NK cells **L.** CD8^+^ T-cells present in subcutaneous (subcut) and liver metastases treated with isotype control (iso) or anti-PD-1 (n=6 experiments). M. Effector CD44^+^ CD62L^-^ T-cells present as a percentage of total CD8^+^ T-cells present in subcutaneous (subcut) and liver metastases treated with isotype control (iso) or anti-PD-1 (n=6 experiments). N. Absolute count of T_REG_ present in subcutaneous (subcut) and liver metastases treated with isotype control (iso) or anti-PD-1 (n=6 experiments). **O-Q.** Mean Florescence intensity (MFI) of **O.** ICOS **P.** CTLA4 and **Q.** CD69 expressed by T_REG_ in subcutaneous (subcut) and liver metastases treated with isotype control (iso) or anti-PD-1 (n=6 experiments). **R&S.** Absolute count of Ly6G+Ly6C^low^ neutrophils present in non-involved liver (LM present) compared to naïve liver of **R.** 4T1 breast cancer **S.** CT26 colorectal cancer (n=2 experiments). K-means clustering to define similar regions using imaging mass cytometry (IMC) within the tissue with the cell density of different cell types shown in each cluster. **T.** Sequential gating strategy for human neutrophil identification (CD11b^hi^CD33^+^HLA-DR^−^CD14^−^CD16^+^) using flow cytometry from freshly isolated lymphocytes. **U.** K-means clustering of 10-cell neighbourhoods to define similar regions using imaging mass cytometry (IMC) fold change within the subcutaneous metastasis, liver metastasis, naïve liver and non-involved liver with the cell density of different cell types shown in each cluster depending on tissue. **V.** Representative images of subcutaneous metastasis, liver metastasis, naïve liver and non-involved liver (LM present) spatially depicting different k-means defined clusters within the tissue. **X.** Comparison of the percentage of different clusters (1-4) defined using k-means clustering observed in the subcutaneous metastasis (n=24 tissue micro-arrays [TMA]), liver metastasis (n=13 TMA), naïve liver (n=4 TMA) and non-involved liver (LM present n=8 TMA). **Y.** Proportional compositions of neighbourhoods (n=10 nearest neighbours) of each seed cell type per tissue type (subcutaneous metastasis, liver metastasis, naïve liver and non-involved liver). Error bars represent mean ±S.E.M. Mann-Whitney test used for comparison of 2 groups (R&S). Kruskal-Wallis test used for comparison of >2 groups (K-Q).

**Supplementary Figure 3.**
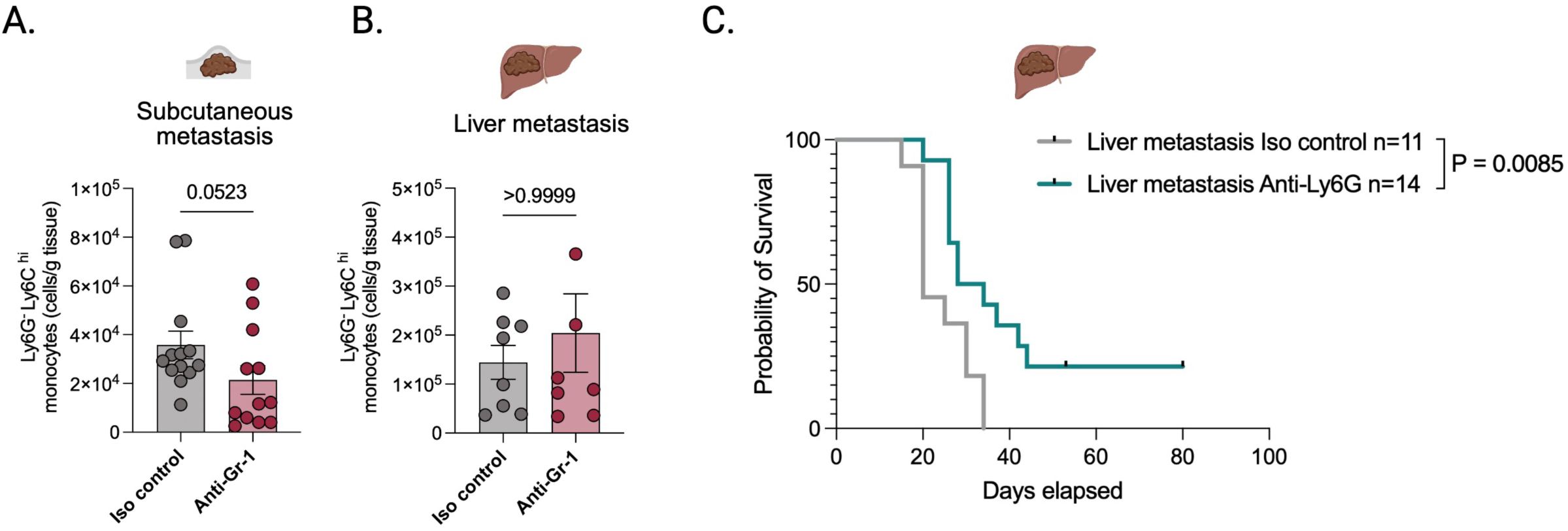
A&B. Infiltration of monocytes is similar in metastases when treated with anti-Gr-1. Absolute count of Ly6G-Ly6C^hi^ monocytes present in **A.** subcutaneous and **B.** liver metastases treated with isotype control or anti-Gr-1 (n=2 experiments, error bars represent median ±S.E.M, Mann Whitney test). **C.** Kaplan Meier survival estimate of mice with liver metastases treated with isotype control or anti-Ly6G. Differences in survival determined by log-rank test.

**Supplementary Figure 4.**
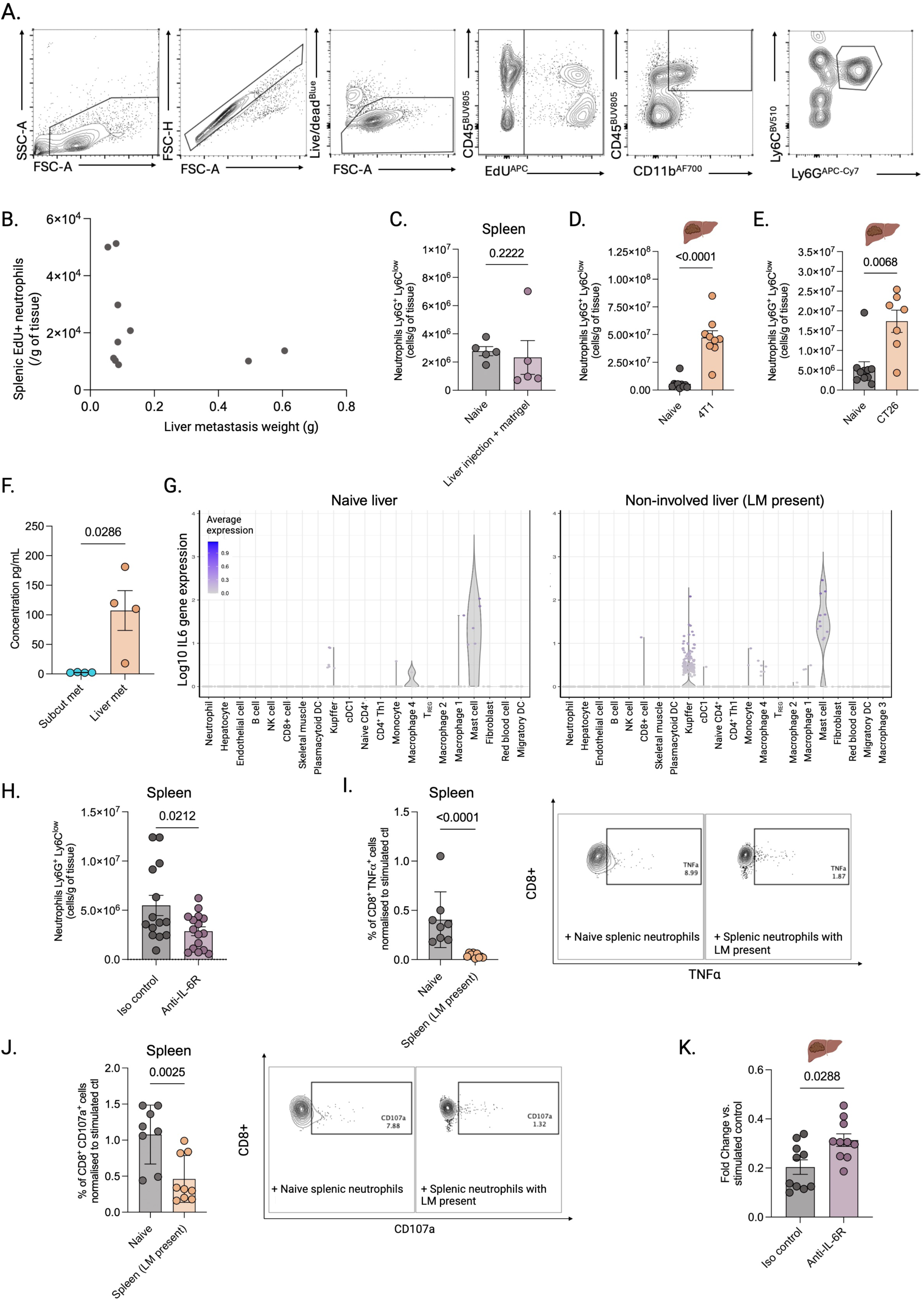
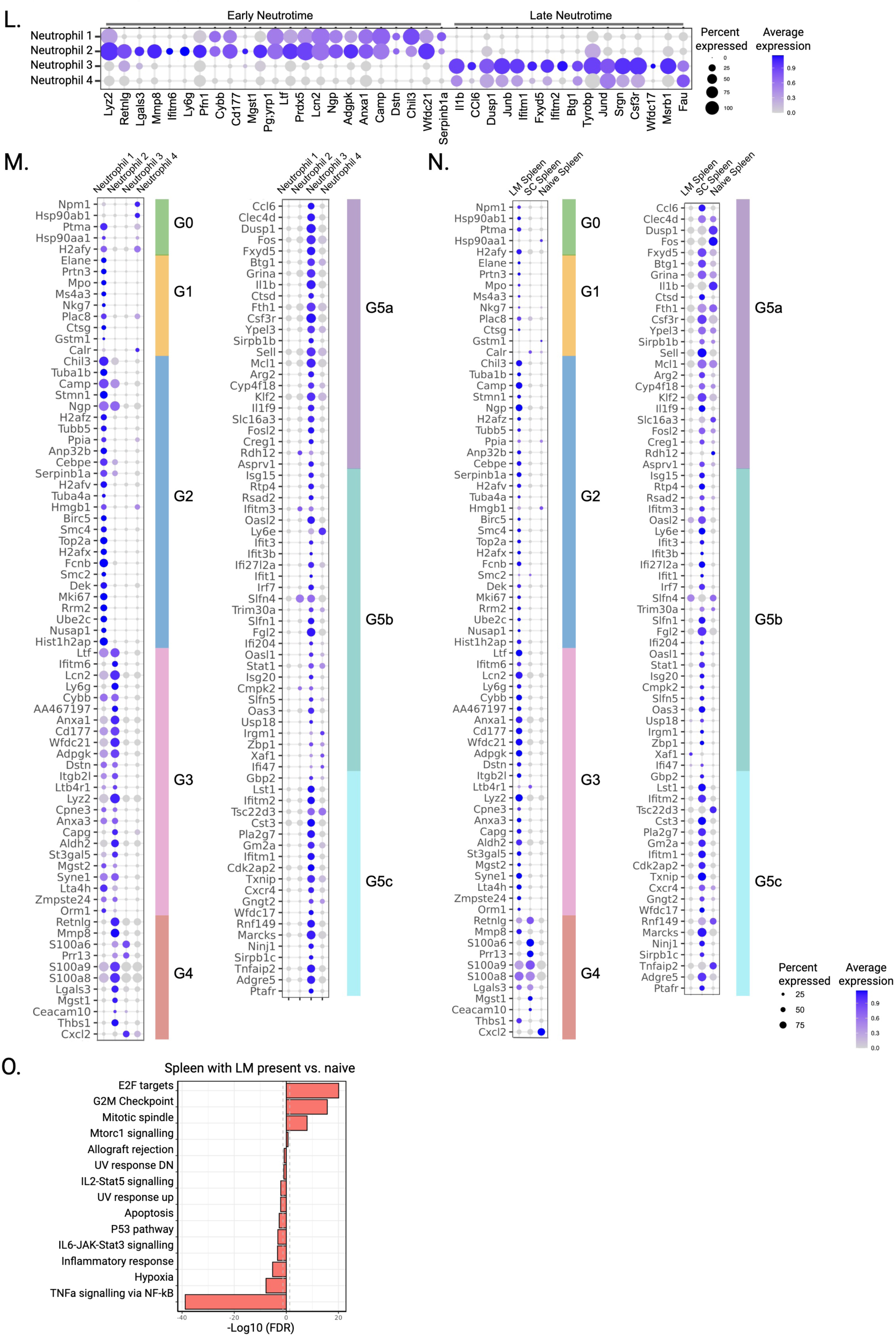
A. Sequential gating strategy to define EdU^+^ mouse neutrophils (EdU^+^ CD45^+^ CD11b^+^ Ly6G^+^ Ly6C^low^) from freshly isolated lymphocytes. **B.** Comparison of absolute count of EdU^+^ splenic Ly6G+Ly6C^low^ neutrophils in spleens vs. weight of liver metastasis. **C.** Absolute count of splenic Ly6G^+^Ly6C^low^ neutrophils of mice with Matrigel injected into the liver compared to naïve livers (n=5 biological replicates). **D&E.** Absolute count of Ly6G^+^Ly6C^low^ neutrophils present in spleens of mice with **D.** 4T1 breast cancer **E.** CT26 colorectal liver metastases present compared to naïve spleens (n=2 experiments). **F.** Concentration of IL6 measured using Luminex immunoassay (n=4 biological replicates per group). **G.** Single cell RNA sequencing of non-involved liver (LM present) compared to naïve liver shows an increase in IL6 gene expression in the Kupffer cell cluster when liver metastases are present. **H.** Absolute count of splenic Ly6G^+^Ly6C^low^ neutrophils of mice with liver metastases present treated with iso control or anti-IL6R (n=2 experiments). **I&J.** Increase in **I.** TNFa and **J.** CD107a expression in CD3/CD28 bead-stimulated splenic T-cells co-cultured 1:1 for 24 hours with neutrophils from spleens of naïve mice or those with a liver metastasis present (n=2 experiments). **K.** Fold change in T cell expansion as measured by cell trace violet dilution after CD3/CD28 bead stimulation following 5-7 days of co-culture 1:1 with neutrophils isolated from spleens of mice with a LM present treated ± anti-IL6R (n=2 experiments). **L.** Dot plot showing early vs. late neutrotime signatures in neutrophil clusters (1-4) in single cell RNA sequencing of splenic neutrophils. Dot size = frequency of cells expressing, colour = average gene expression. **M.** Dot plot showing Xie G0-G5 signatures in neutrophil clusters (1-4) in single cell RNA sequencing of splenic neutrophils. Dot size = frequency of cells expressing, colour = average gene expression. **N.** Hallmark pathway analysis of differentially expressed genes using single cell RNA sequencing of splenic neutrophils when a LM is present compared to naïve spleen. **O.** Dot plot showing Xie G0-G5 signatures in single cell RNA sequencing of splenic neutrophils in naïve spleens or spleens with a subcutaneous or liver metastasis present. Dot size = frequency of cells expressing, colour = average gene expression. Error bars represent mean ±S.E.M. Mann Whitney test used for all statistical analyses.

**Supplementary Figure 5.**
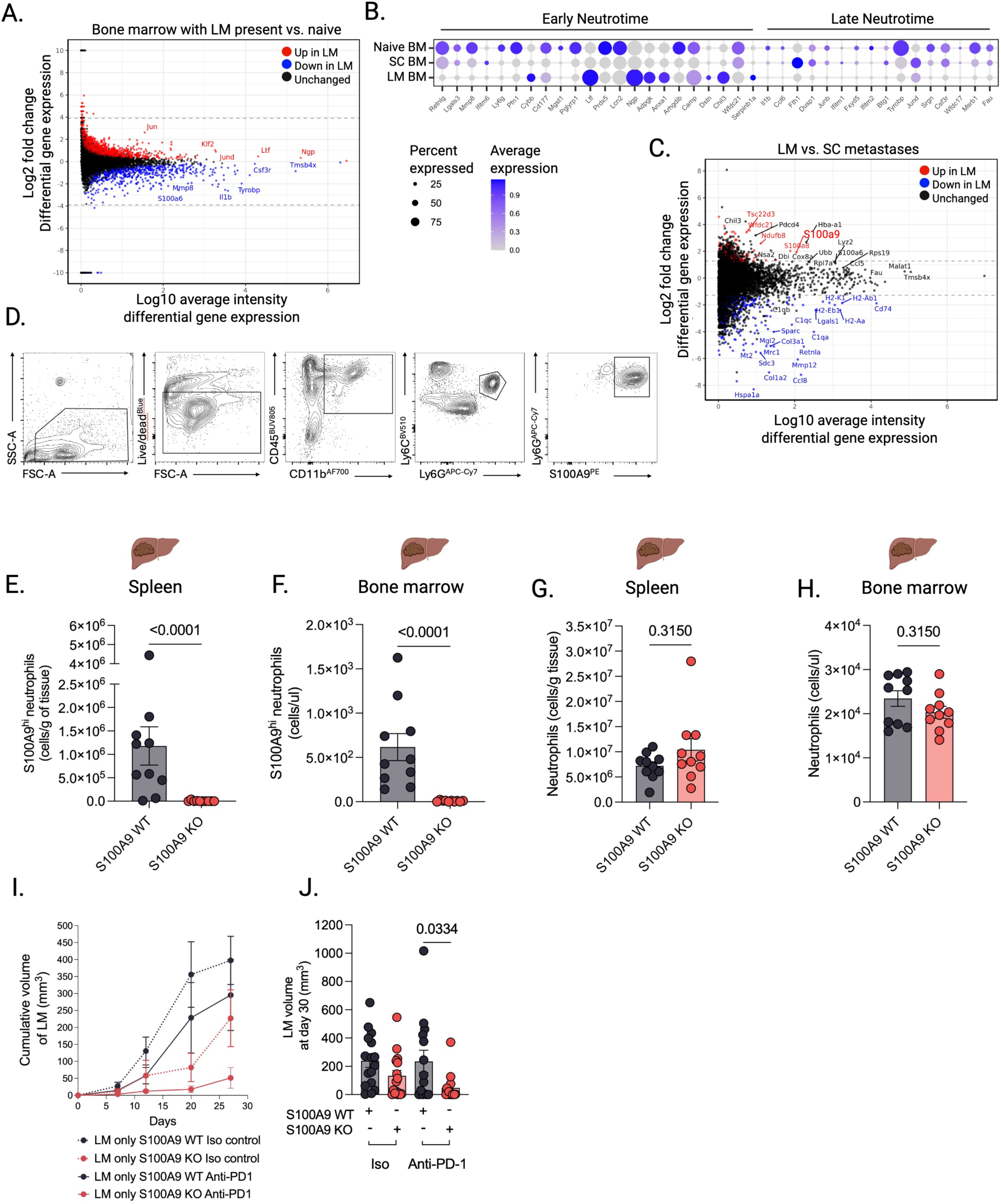
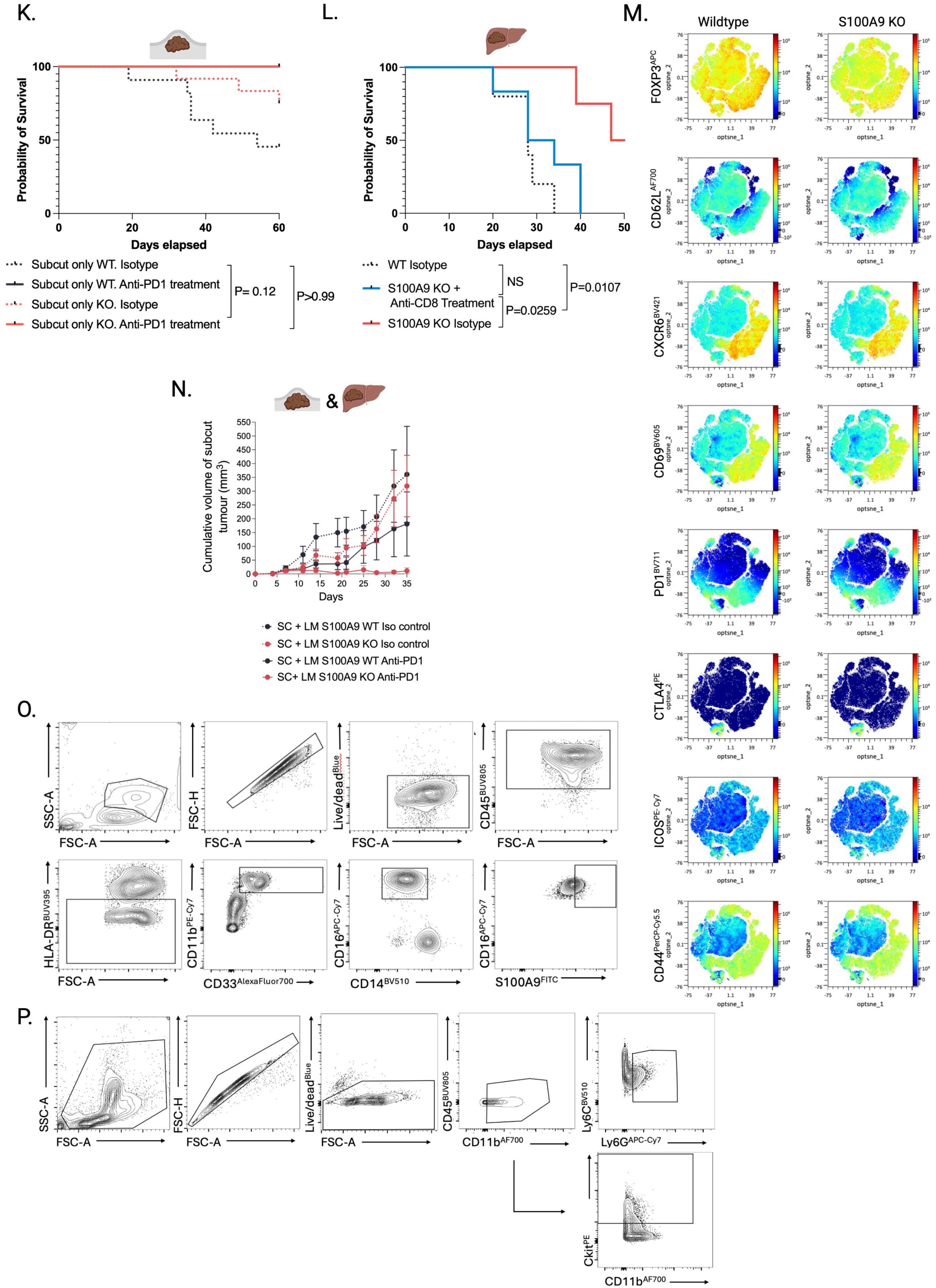
**B.** MA plot showing differential gene expression of single cell sequenced bone marrow neutrophils in mice bearing liver metastases compared to normal bone marrow. **C.** Single cell analysis dot plots comparing gene expression of early and late neutrotime signatures of neutrophils in normal bone marrow or bone marrow (BM) of mice with a subcutaneous (SC) or liver metastasis (LM) present. **D.** MA plot depicting differential gene expression of single cell sequenced neutrophils infiltrating liver vs. subcutaneous metastases. **E.** Sequential gating strategy to define S100A9^hi^ mouse neutrophils (S100A9^hi^ CD45^+^ CD11b^+^ Ly6G^+^ Ly6C^low^) from freshly isolated lymphocytes. **E&F.** Absolute count of S100A9^hi^ Ly6G^+^ Ly6C^low^ neutrophils present in **E.** spleens and **F.** bone marrow of S100A9 knockout (KO) vs. wildtype (WT) mice bearing liver metastases (n=2 experiments) **G&H.** Absolute count of Ly6G^+^ Ly6C^low^ neutrophils present in spleens **G.** and bone marrow **H.** of S100A9 knockout (KO) vs. wildtype (WT) mice bearing liver metastases (n=2 experiments) **I.** Photographs taken of liver metastases in S100A9 knockout (KO) vs. wildtype (WT) at autopsy performed on day 14 post 4434 cell injection. **J.** Growth of liver metastases in S100A9 knockout (KO) vs. wildtype (WT) mice measured using ultrasound treated with isotype control (iso) or anti-PD-1 (n=5 biological replicates). **K.** Volume of liver metastases (mm^3^) at day 30 in S100A9 knockout (KO) vs. wildtype (WT) mice measured using ultrasound treated with isotype control (iso) or anti-PD-1. **L.** Kaplan Meier survival estimate of wildtype (black) or S100A9 KO mice (orange) with subcutaneous metastases treated with isotype control (iso, dashed line) or anti-PD-1 (continuous line). Differences in survival determined by log-rank test. **M.** Kaplan Meier survival estimate of S100A9 KO vs. WT mice with liver metastases ± CD8^+^ cell depletion. Differences in survival determined by log-rank test. **N.** Opt-SNE plots of marker expression on CD4^+^ T cells in non-involved livers of mice with S100A9 KO vs. wildtype culled at day 14 post injection.

## Author contributions

Conceptualization RL, ES, MM, LP, MD Data curation RL, SH, TMS, LP, MD, NR, ZR, CB, SB Formal analysis RL, LP, MD, TMS, NR, ZR, SB, OK Funding acquisition RL, ES, MM, PL Investigation RL, SH, TMS, LP, MD, NR, ZR, SB, OK, CB, GAM, SM, SB, JD, AM, GP, AC, VP, IM Methodology RL, SH, TMS, LP, MD, NR, ZR, CB, VP, IM Project administration RL, ES, MM, PL Resources RL, ES, MM, PL, PS, AG, BD, JP, VP, IM, GP Supervision ES, MM, PL Validation RL, SB Visualization RL, SB, ES Writing – original draft RL, ES, MM Writing – review & editing all

## Acknowledgements

We acknowledge the Biological Research Facility including Lauren Chisholm, Michael Nagliati, Ryan Hoskins, Nicholas Chisholm, Sam Cooper and Rekha Subramaniam for animal husbandry and procedures, the Genomics Science Technology Platform, particularly Hubert Slawinski, and Marg Crawford, for their contributions to the single-cell capture experiment, library construction, and sequencing, the Experimental Histology Platform especially Nisha Bhardwaj, Ania Mikolajczak, Richard Stone and Emma Nye, the Flow Cytometry Platform particularly Gemma Foulds and Philip Hobson, the In Vivo Imaging platform especially Peter Johnson and the Cell Science Platform all at The Francis Crick Institute, London. The Flow Cytometry Facility particularly Janani Sivakumaran-Nguyen and Sam Blanchett at The Pears Building, Institute of Immunity and Transplantation. We are very grateful to all patients, healthy volunteers and their families who participated in this study and to all clinical staff who helped with participant recruitment, including the Tissue Access for Patient Benefit project at The Royal Free Hospital and the melanoma research team at The Christie Hospital in Manchester. Figures made using BioRender https://BioRender.com.

## References

1. Schmid, P. et al. Pembrolizumab for Early Triple-Negative Breast Cancer. New England Journal of Medicine 382, 810–821 (2020).

2. André, T. et al. Pembrolizumab in Microsatellite-Instability–High Advanced Colorectal Cancer. New England Journal of Medicine 383, 2207–2218 (2020).

3. Tawbi, H. A. et al. Relatlimab and Nivolumab versus Nivolumab in Untreated Advanced Melanoma. New England Journal of Medicine 386, 24–34 (2022).

4. Wolchok, J. D. et al. Final, 10-Year Outcomes with Nivolumab plus Ipilimumab in Advanced Melanoma. New England Journal of Medicine 392, 11–22 (2025).

5. Sharma, P. et al. Immune checkpoint therapy—current perspectives and future directions. Cell 186, 1652–1669 (2023).

6. Cogdill, A. P., Andrews, M. C. & Wargo, J. A. Hallmarks of response to immune checkpoint blockade. British Journal of Cancer 2017 117:1 117, 1–7 (2017).

7. Yoo, S. K. et al. Prediction of checkpoint inhibitor immunotherapy efficacy for cancer using routine blood tests and clinical data. Nature Medicine 2025 31:3 31, 869–880 (2025).

8. Topalian, S. L. et al. Five-Year Survival and Correlates Among Patients With Advanced Melanoma, Renal Cell Carcinoma, or Non–Small Cell Lung Cancer Treated With Nivolumab. JAMA Oncol 5, 1411–1420 (2019).

9. Pires da Silva, I., et al. Site-specific response patterns, pseudoprogression, and acquired resistance in patients with melanoma treated with ipilimumab combined with anti–PD-1 therapy. Cancer 126, 86–97 (2020).

10. Taner, T. et al. Donor-specific hypo-responsiveness occurs in simultaneous liver-kidney transplant recipients after the first year. Kidney Int 93, 1465–1474 (2018).

11. Calne, R. Y. et al. Induction of immunological tolerance by porcine liver allografts. Nature 223, 472–476 (1969).

12. Lee, J. C., et al. Regulatory T cell control of systemic immunity and immunotherapy response in liver metastasis. *Sci Immunol* 5, (2020).

13. McFarlane, A. J., Fercoq, F., Coffelt, S. B. & Carlin, L. M. Neutrophil dynamics in the tumor microenvironment. J Clin Invest 131, (2021).

14. Coffelt, S. B., Wellenstein, M. D. & De Visser, K. E. Neutrophils in cancer: neutral no more. Nature Reviews Cancer 2016 16:7 16, 431–446 (2016).

15. Geh, D. et al. Neutrophils as potential therapeutic targets in hepatocellular carcinoma. Nature Reviews Gastroenterology & Hepatology 2022 19:4 19, 257–273 (2022).

16. Blattner, C. et al. CCR5+ myeloid-derived suppressor cells are enriched and activated in melanoma lesions. Cancer Res 78, 157–167 (2018).

17. Canè, S. et al. Neutralization of NET-associated human ARG1 enhances cancer immunotherapy. Sci Transl Med 15, (2023).

18. Veglia, F. et al. Fatty acid transport protein 2 reprograms neutrophils in cancer. Nature 2019 569:7754 569, 73–78 (2019).

19. Pallett, L. J. et al. Metabolic regulation of hepatitis B immunopathology by myeloid-derived suppressor cells. Nat Med 21, 591 (2015).

20. Yu, J. et al. Liver metastasis restrains immunotherapy efficacy via macrophage-mediated T cell elimination. Nat Med 27, 152–164 (2021).

21. Cassidy, M. R. et al. Neutrophil to Lymphocyte Ratio is Associated With Outcome During Ipilimumab Treatment. EBioMedicine 18, 56–61 (2017).

22. Capone, M. et al. Baseline neutrophil-to-lymphocyte ratio (NLR) and derived NLR could predict overall survival in patients with advanced melanoma treated with nivolumab. J Immunother Cancer 6, (2018).

23. Valero, C. et al. Pretreatment neutrophil-to-lymphocyte ratio and mutational burden as biomarkers of tumor response to immune checkpoint inhibitors. Nature Communications 2021 12:1 12, 1–9 (2021).

24. Yang, L. et al. DNA of neutrophil extracellular traps promotes cancer metastasis via CCDC25. Nature 2020 583:7814 583, 133–138 (2020).

25. Hsu, B. E. et al. Immature Low-Density Neutrophils Exhibit Metabolic Flexibility that Facilitates Breast Cancer Liver Metastasis. Cell Rep 27, 3902–3915.e6 (2019).

26. Grieshaber-Bouyer, R. et al. The neutrotime transcriptional signature defines a single continuum of neutrophils across biological compartments. Nat Commun 12, (2021).

27. Xie, X. et al. Single-cell transcriptome profiling reveals neutrophil heterogeneity in homeostasis and infection. Nat Immunol 21, 1119–1133 (2020).

28. Evrard, M. et al. Developmental Analysis of Bone Marrow Neutrophils Reveals Populations Specialized in Expansion, Trafficking, and Effector Functions. Immunity 48, 364–379.e8 (2018).

29. Hobbs, J. A. R. et al. Myeloid Cell Function in MRP-14 (S100A9) Null Mice. Mol Cell Biol 23, 2564–2576 (2003).

30. Nacken, W. & Kerkhoff, C. The hetero-oligomeric complex of the S100A8/S100A9 protein is extremely protease resistant. FEBS Lett 581, 5127–5130 (2007).

31. Bollyky, P. L. et al. CD44 Costimulation Promotes FoxP3+ Regulatory T Cell Persistence and Function via Production of IL-2, IL-10, and TGF-β. The Journal of Immunology 183, 2232–2241 (2009).

32. Burton, O. T. et al. The tissue-resident regulatory T cell pool is shaped by transient multi-tissue migration and a conserved residency program. Immunity 57, 1586–1602.e10 (2024).

33. Quail, D. F. et al. Neutrophil phenotypes and functions in cancer: A consensus statement. Journal of Experimental Medicine 219, 39 (2022).

34. Hedrick, C. C. & Malanchi, I. Neutrophils in cancer: heterogeneous and multifaceted. Nature Reviews Immunology 2021 22:3 22, 173–187 (2021).

35. Mackey, J. B. G. et al. Maturation, developmental site, and pathology dictate murine neutrophil function 1. doi:10.1101/2021.07.21.453108.

36. Benguigui, M. et al. Interferon-stimulated neutrophils as a predictor of immunotherapy response. Cancer Cell 42, 253–265.e12 (2024).

37. Hsu, B. E. et al. Immature Low-Density Neutrophils Exhibit Metabolic Flexibility that Facilitates Breast Cancer Liver Metastasis. Cell Rep 27, 3902–3915.e6 (2019).

38. Leslie, J. et al. CXCR2 inhibition enables NASH-HCC immunotherapy. Gut 71, 2093–2106 (2022).

39. Wagner, N. B. et al. Tumor microenvironment-derived S100A8/A9 is a novel prognostic biomarker for advanced melanoma patients and during immunotherapy with anti-PD-1 antibodies. J Immunother Cancer 7, 343 (2019).

40. Cheng, P. et al. Inhibition of dendritic cell differentiation and accumulation of myeloid-derived suppressor cells in cancer is regulated by S100A9 protein. J Exp Med 205, 2235 (2008).

41. Sinha, P. et al. Proinflammatory S100 Proteins Regulate the Accumulation of Myeloid-Derived Suppressor Cells. The Journal of Immunology 181, 4666–4675 (2008).

42. Wang, Q. et al. S100A8/A9: An emerging player in sepsis and sepsis-induced organ injury. Biomedicine & Pharmacotherapy 168, 115674 (2023).

43. Sprenkeler, E. G. G. et al. S100A8/A9 Is a Marker for the Release of Neutrophil Extracellular Traps and Induces Neutrophil Activation. Cells 2022, Vol. 11, Page 236 11, 236 (2022).

44. Schenten, V. et al. Secretion of the phosphorylated form of S100A9 from neutrophils is essential for the proinflammatory functions of extracellular S100A8/A9. Front Immunol 9, 294341 (2018).

45. Pruenster, M. et al. E-selectin-mediated rapid NLRP3 inflammasome activation regulates S100A8/S100A9 release from neutrophils via transient gasdermin D pore formation. Nat Immunol 24, 2021–2031 (2023).

46. Manitz, M.-P. et al. Loss of S100A9 (MRP14) Results in Reduced Interleukin-8-Induced CD11b Surface Expression, a Polarized Microfilament System, and Diminished Responsiveness to Chemoattractants In Vitro. Mol Cell Biol 23, 1034–1043 (2003).

47. Kucykowicz, S. et al. Isolation of human intrahepatic leukocytes for phenotypic and functional characterization by flow cytometry. STAR Protoc 3, (2022).

48. Hao, Y. et al. Dictionary learning for integrative, multimodal and scalable single-cell analysis. Nat Biotechnol 42, 293–304 (2024).

49. Ahlmann-Eltze, C. & Huber, W. glmGamPoi: fitting Gamma-Poisson generalized linear models on single cell count data. Bioinformatics 36, 5701–5702 (2021).

50. Magness, A. et al. Deep cell phenotyping and spatial analysis of multiplexed imaging with TRACERx-PHLEX. Nature Communications 2024 15:1 15, 1–20 (2024).

51. Giangreco, G. & Sahai, E. STC2 coordinates multicellular Ca2+ waves in stromal fibroblasts leading to immune exclusion. In submission (2025).

52. R Foundation for Statistical Computing. R: A Language and Environment for Statistical Computing. (2024).

53. Schneider, C. A., Rasband, W. S. & Eliceiri, K. W. NIH Image to ImageJ: 25 years of image analysis. Nat Methods 9, 671–675 (2012).

