## Supplementary tables for "Splenic granulopoiesis and S100A9 drive resistance to checkpoint inhibitors conferred by liver metastases"

**Supplementary Table 1. Baseline characteristics**

|  | **All** | | | **Melanoma** | | **NSCLC** | |
| --- | --- | --- | --- | --- | --- | --- | --- |
|  | **Total** | **No liver mets** | **Liver mets** | **Melanoma and no liver mets** | **Melanoma and liver mets** | **NSCLC and no liver mets** | **NSCLC and liver mets** |
| **All** | 1024 | 793 | 231 | 476 | 165 | 317 | 66 |
| **Sex** |  |  |  |  |  |  |  |
| Female | 441 | 342 | 99 | 185 | 70 | 157 | 29 |
| Male | 583 | 451 | 132 | 291 | 95 | 160 | 37 |
| **Age** |  |  |  |  |  |  |  |
| <50 | 92 | 69 | 23 | 61 | 22 | 8 | 1 |
| 50-64 | 652 | 511 | 141 | 288 | 104 | 223 | 37 |
| >65 | 280 | 213 | 67 | 127 | 39 | 86 | 28 |
| **Performance status** |  |  |  |  |  |  |  |
| 0 | 614 | 479 | 135 | 314 | 100 | 165 | 35 |
| 1 | 330 | 253 | 77 | 123 | 50 | 130 | 27 |
| 2+ | 67 | 50 | 17 | 34 | 13 | 16 | 4 |
| Unknown | 13 | 11 | 2 | 5 | 2 | 6 | 66 |
| **Treatment** |  |  |  |  |  |  |  |
| Atezolizumab or pembrolizumab 1st line (advanced)--nsclc | 383 | 317 | 66 | 0 | 0 | 317 | 66 |
| Nivolumab and Ipilimumab (advanced)--Melanoma | 336 | 239 | 97 | 239 | 97 | 0 | 0 |
| Nivolumab/pembrolizumab (advanced)--Melanoma | 305 | 237 | 68 | 237 | 68 | 0 | 0 |
| Other advanced |  |  |  |  |  |  |  |
| **Blood tests** |  |  |  |  |  |  |  |
| Median neutrophils | 5.7 | 5.6 | 6 | 5.1 | 5.25 | 6.7 | 9 |
| Median lymphocytes | 1.45 | 1.5 | 1.4 | 1.5 | 1.5 | 1.4 | 1.4 |
| Median LDH | 256 | 244 | 308 | 239 | 330 | 255 | 283 |
| **BRAF** |  |  |  |  |  |  |  |
| Wildtype | 161 | 82 | 79 | 82 | 79 | 0 | 0 |
| Mutant | 48 | 30 | 18 | 30 | 18 | 0 | 0 |
| Unknown/NA | 815 | 681 | 134 | 364 | 68 | 317 | 66 |

**Supplementary table 2. Overall survival patients with melanoma treated with 1^st^ line CPI**

| **Diagnosis** | **n/events** | **Median OS (95% CI)** | **1-year OS (95% CI)** | **Univariate HR (95% CI)** | **Multivariable HR (95% CI)** |
| --- | --- | --- | --- | --- | --- |
| **No liver metastasis** | 476/240 | 44.9 (32.4-55.7) | 73.1% (69.2%-77.2%) | 1. (ref.) | 1. (ref.) |
| **Liver metastasis** | 165/105 | 15.2 (11.6-24.5) | 56.4% (49.3%-64.5%) | 1.60 (1.27 - 2.01) | 1.69 (1.30 - 2.20) |

Adjusted for age, sex, treatment, BRAF (wildtype/mutant) and performance status

**Supplementary table 3. Overall survival patients with NSCLC treated with 1^st^ line CPI**

| **Diagnosis** | **n/events** | **Median OS months (95% CI)** | **1-year OS (95% CI)** | **Univariate HR (95% CI)** | **Multivariable HR (95% CI)** |
| --- | --- | --- | --- | --- | --- |
| **No liver metastasis** | 317/209 | 20.9 (17.0-25.5) | 62.5% (57.4%-68.0%) | 1. (ref.) | 1. (ref.) |
| **Liver metastasis** | 66/56 | 8.1 (4.8-16.2) | 40.9% (30.6%-54.7%) | 1.74 (1.29 - 2.34) | 1.62 (1.19 - 2.21) |

Adjusted for age, sex, treatment and performance status
